# m^6^A RNA methylation-dependent APC translation is required for local translation of beta-actin and axon development

**DOI:** 10.1101/2020.11.14.382556

**Authors:** Loic Broix, Rohini Roy, Ikumi Oomoto, Hiroki Umeshima, Dan Ohtan Wang

**Affiliations:** RIKEN Center for Biosystems Dynamics Research (BDR), Kobe, Japan; Graduate School of Biostudies, Kyoto University, Kyoto, Japan; Institute for Integrated Cell-Material Sciences (iCeMS), Kyoto University, Kyoto, Japan; New York University (NYU), Abu Dhabi, United Arab Emirates

## Abstract

Subcellular mRNA localization and local translation are crucial for spatially regulated gene expression in neurons. However, the precise mechanisms governing the selective transport and translation of mRNAs contributing to processes such as axonal growth and branching are only partially understood. Here, we present evidence of N6-methyladenosine (m^6^A)-mediated translational control of the RNA-binding protein, APC, influencing cytoskeletal dynamics at the growth cone. Our findings demonstrate that m^6^A modifications occur on *Apc* mRNA, facilitating its recognition by YTHDF1 to promote APC translation in neuronal somata. Additionally, we observe that disrupting the m^6^A pathway impairs the transport and local translation of β-actin mRNA in the axon and growth cones, a deficiency that can be rescued by the exogenous expression of APC protein in cultured neurons. Furthermore, we establish the essential role of YTHDF1 in axon development, particularly in callosal projection neurons during cortical development. Our findings suggest a novel mechanism involving m^6^A-mediated regulation of APC protein translation, linking epitranscriptomics to axonal mRNA targeting, cytoskeletal dynamics, and axon development.

## Introduction

Subcellular mRNA localization and local translation serve as a crucial mechanism for spatially regulated gene expression within cells. This phenomenon is particularly prominent in highly polarized cells such as neurons, where a multitude of mRNAs can be directed to distant compartments of axons and dendrites (Holt and Schuman, 2019; Bourke et al., 2023). In these specialized regions, regulated local translation contributes to the synthesis of proteins essential for processes such as axonal and dendritic growth, branching and maintenance, and synaptic plasticity (Batista and Hengst, 2016). In particular, local translation at the axonal growth cone is essential for growth cone guidance, mediating rapid cytoskeletal remodeling in response to environmental cues. The existing model for the intracellular distribution of mRNAs in neurons primarily involves their integration into membrane-less ribonucleoprotein (RNP) granules and association with molecular motors for their active long-distance transport along microtubules (MTs) into distal parts of dendrites or axons (Broix et al., 2019; Dalla Costa et al., 2021). RNP granules are formed through multivalent interactions between mRNAs containing cis-acting recognition motifs, structural elements typically located in their 3′ UTRs, and RNA-binding proteins (RBPs) (Tushev et al., 2018).

While widely recognized as a tumor suppressor, Adenomatous polyposis coli (APC) protein has also been identified as an RBP in migrating fibroblasts, *Drosophila* oocytes and neurons (Mili et al., 2008; Preitner et al., 2014; Vaishali et al., 2020). APC binds to the 3’UTR of *β2B-tubulin* mRNA, among 260 other target mRNAs encoding mainly cytoskeletal regulators, and promotes its axonal localization and translation, which ultimately regulates dynamic microtubules in the growth cone (Preitner et al., 2014). Furthermore, recent work has demonstrated that APC serves as the principal regulatory element in an *in vitro* reconstituted mammalian anterograde-directed RNP complex. It achieves this by binding to guanine-rich sequences in the 3’ UTR of β-actin and β2B-tubulin mRNAs, subsequently linking them to kinesin-2 to facilitate the recruitment of the complex onto microtubules for their transport (Baumann et al., 2020). Beyond its role as an RBP, APC exhibits direct interaction with the microtubule lattice and can bind to growing microtubule ends through its interaction with end-binding (EB) proteins, which are a core family of MT plus-end tracking proteins that have profound effects on the shape of the microtubule network (Nakamura et al., 2001; Zumbrunn et al., 2001). Moreover, APC can bind actin and nucleate unbranched actin filaments either independently or in collaboration with formins (Moseley et al., 2007; Okada et al., 2010; Breitsprecher et al., 2012). Consistent with the above roles of APC in cytoskeleton regulation, APC was shown to be crucial for cell motility and polarity, as well as neurite outgrowth and has been described as a risk gene in neurological diseases such as autism and schizophrenia (Barber et al., 1994; Raedle et al., 2001; Watanabe et al., 2004; Zhou et al., 2004; Cui et al., 2005; Finch et al., 2005; Heald et al., 2007; Zhou et al., 2007; Yokota et al., 2009; Mohn et al., 2014).

To date, no universal localization cis-acting elements have been identified among all distally-localized mRNAs which implies that there might be additional mechanisms beyond the primary sequences of the RNA to provide localization information in neurons. Epitranscriptomics, a field concerning dynamic genetic deciphering based on type- and site-specific RNA modifications, has recently emerged as a versatile and powerful post-transcriptional regulatory pathway in the central nervous system (Wang, 2020; Madugalle et al., 2020). N6-methyladenosine (m^6^A) is the best characterized and the most abundant internal mRNA modification in mammals which decorates up to half of the entire poly(A) transcriptome in the brain tissue (Livneh et al., 2020). Accordingly, m^6^A has been shown to play critical roles in embryonic neurogenesis and differentiation (Yoon et al., 2017; Li et al., 2018; Wang et al., 2018; Edens et al., 2019), axon guidance and regeneration (Yu et al., 2018; Weng et al., 2018; Zhuang et al., 2019; Yu et al., 2021; Worpenberg et al., 2021; Huang et al., 2022; Wang et al., 2023), synaptic function (Merkurjev et al., 2018; Martinez De La Cruz et al. 2021), as well as learning and memory (Hess et al., 2013; Widagdo et al., 2016; Walters et al., 2017; Shi et al., 2018; Zhuang et al., 2023). Our advanced understanding of m^6^A can be attributed to the identification of its “writer” complex (containing METTL3, METTL14, WTAP, and other subunits of the methyltransferase complex), “erasers” (FTO and ALKBH5), and “readers” (YTH family proteins, hnRNPC, hnRNPG, IGF2BPs, eIF3, etc.).The recognition and binding of the YTHDF reader proteins to m^6^A-modified mRNAs has been shown to regulate multiple facets of gene expression such as mRNA nuclear export, stability, translation and phase separation (Roundtree et al., 2017; Edens et al., 2019; Wang et al., 2014; Zaccara and Jaffrey 2020; Meyer et al., 2015; Coots et al., 2017; Wang et al., 2015; Ries et al., 2019; Fu et al., 2020). We have previously reported that a subset of hypermethylated transcripts coding for synapse-related proteins are enriched in the synaptic compartments, suggesting that m^6^A modification could have a function in the subcellular localization and local translation of specific mRNAs (Merkurjev et al., 2018). This is consistent with a recent study that identified a subset of learning-related m^6^A-modified RNAs enriched at synapses essential for the consolidation of fear-extinction memory (Madugalle et al., 2023). Nevertheless, two recent studies have found conflicting results regarding the involvement of m^6^A signaling in controlling mRNA localization in neurons. Flamand and Meyer reported that m^6^A methylation in the 3’UTR of hundreds of mRNAs facilitates their distal transport to neurites (Flamand and Meyer, 2022), whereas Loedige et al. described that m^6^A-modified transcripts are predominantly restricted to neuronal cell soma due to their reduced stability (Loedige et al., 2023). Further mechanistic investigations are thus necessary for understanding how m^6^A modification and its associated machinery control mRNA localization and local translation in neurons. A cell biological functional link between the m^6^A machinery and the modulation of the cytoskeleton, a process crucial for axonal development and engages local protein synthesis, could provide insights into the potential functional mechanism of m^6^A.

Here, we provide evidence of m^6^A-mediated epitranscriptomic regulation of APC translation that modulates cytoskeleton dynamics at the axonal growth cones. We report that m^6^A marks are installed on *Apc* mRNA and that the subsequent recognition and binding by YTHDF1 regulates APC translation within neuronal somata. We further show that interfering m^6^A pathway disrupts the transport and local translation of β-actin mRNA in the axon and growth cone, which can be rescued by exogenously expressed APC protein. We validate the functional requirement of YTHDF1 for commissural axon development in the corpus callosum during cortical development. Together, our data reveal that the m^6^A-mediated control of APC translation in the cell soma modulates β-actin axonal mRNA trafficking and local translation that are required for the progression of axon development.

## Results

### m^6^A modification is installed on *Apc* mRNA but does not impact *Apc* mRNA stability and nuclear export in neurons

While previous studies, including our own, utilizing high-throughput antibody-based technologies indicated that *Apc* mRNA is likely to be m^6^A-modified (Meyer et al., 2012; Ke et al., 2015; Wang et al., 2015; Shi et al., 2018; Merkurjev et al., 2018), the predicted sites of modification have not been validated. We employed an antibody-independent m^6^A detection method named single-base elongation- and ligation-based qPCR amplification (SELECT) to confirm selective m^6^A marks on *Apc* mRNA (Xiao et al., 2018). Three adenosines in mouse *Apc* mRNA (NM_007462.3) were chosen for validation, with one in the coding sequence (A7836) and two in the 3’UTR (A11110, A12324), as predicted by high-throughput analysis (**Fig. 1A**). Neuro2a cells were transfected with either a plasmid overexpressing the mouse ALKBH5 m^6^A-demethylase protein or an Empty vector. qPCR amplification for the three selected sites relative to a control adenosine site (lacking a canonical m^6^A motif) revealed a significant increase in relative ligation products for all three sites in the cells transfected with the ALKBH5-encoding plasmid compared to the control, confirming these three adenosines as m^6^A-methylated sites in *Apc* mRNA (**Fig. 1B and 1C**). It is noteworthy that overexpression of ALKBH5 did not affect *Apc* mRNA abundance. Given the well-established role of m^6^A addition and the downstream reading of the m^6^A signal in mRNA regulation, we sought to investigate the effect of YTHDF1 and METTL14 knockdowns on *Apc* mRNA level and localization in cultured neurons. The consensus is that YTHDF1 regulates translation, YTHDF2 facilitates degradation and YTHDF3 exerts its function through interactions with either YTHDF1 or YTHDF2 (He and He, 2021). As we did not observe any alteration in *Apc* mRNA expression following ALKBH5 overexpression, we selected YTHDF1 among the three YTHDF proteins for further investigation. METTL14 was selected over METTL3 due to the high level of toxicity observed in neuronal cells following METTL3 knockdown. Firstly, shRNAs against *Ythdf1* and *Mettl14* were validated and showed significant reductions of *Ythdf1* and *Mettl14* mRNA expressions by qPCR and YTHDF1 and METTL14 protein levels by immunostainings on DIV6 neurons transfected at DIV2 (**Fig. S1**). Subsequently, qPCR analysis from Neuro2a cells transfected with either sh*Ythdf1* or sh*Mettl14* showed no difference in the expression of *Apc* mRNA in comparison to the control (**Fig. 1D**). We then utilized the commercially available ViewRNA ISH cell assay kit to detect single molecule *Apc* mRNA with high sensitivity in cultured DIV6 neurons, and we observed that YTHDF1 or METTL14 downregulations had no impact on the total level of *Apc* mRNAs in the cell soma as well as in the nucleus/cytosol ratio (**Fig. 1E and 1F**). Overall, this data indicates that *Apc* mRNA undergoes m^6^A modification, but it is not essential for the expression, stability, and nuclear export of *Apc* mRNA in neurons.

**Figure 1.**
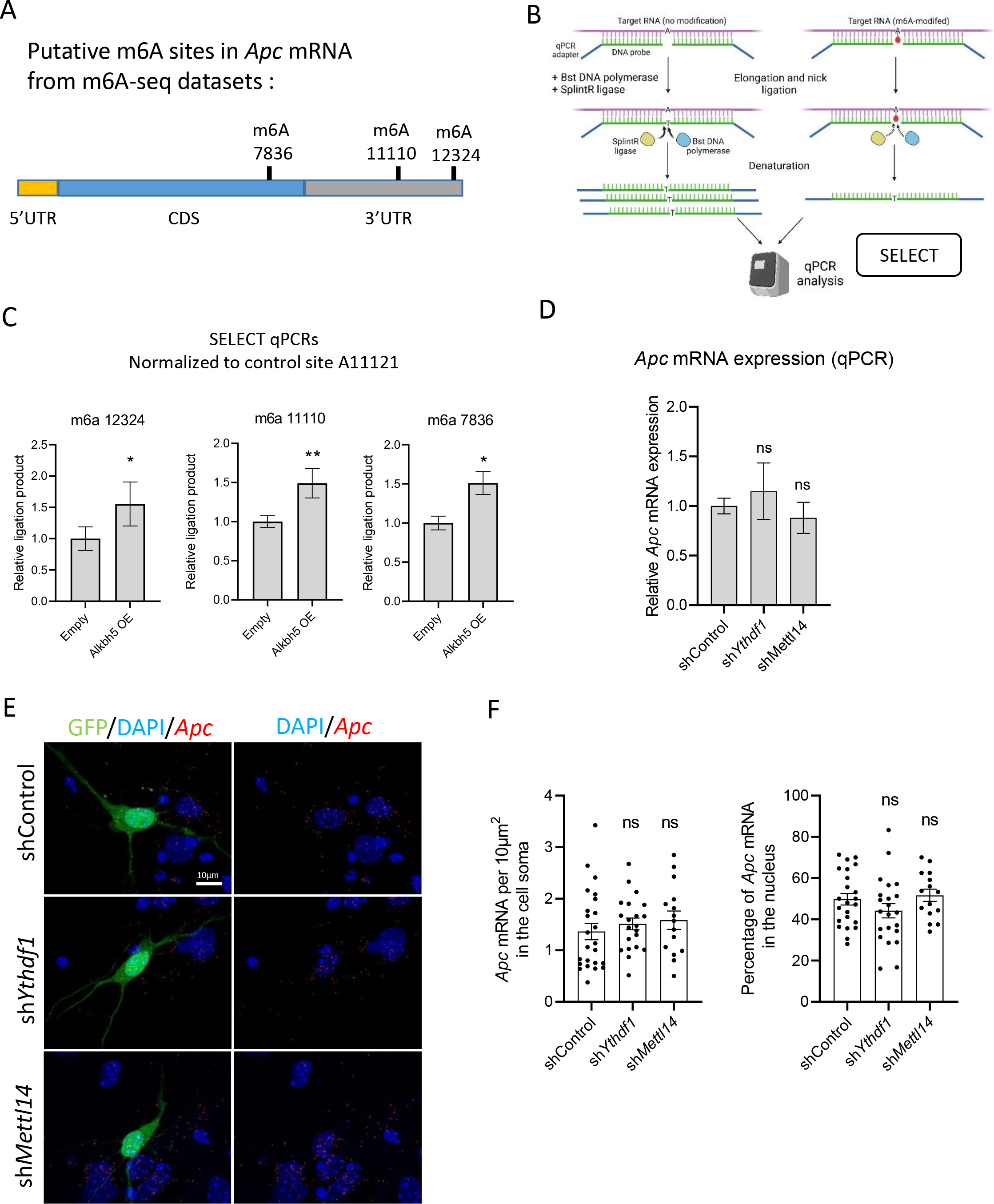
*Apc* mRNA is m^6^A-methylated but m^6^A machinery deregulation does not impact its expression nor nuclear export. (A) Linear schematic representation of *Apc* mRNA (5’UTR in yellow; CDS in blue; 3’UTR in gray) indicating the positions of three putative m^6^A-modified nucleotides selected for further validation by SELECT. (B) Illustration of the SELECT m^6^A detection method. (C) Measurement of m^6^A level by SELECT on A7836, A11110 and A12324 of *Apc* mRNA in Neuro2a cells transfected with an empty vector or an ALKBH5-expressing vector. Ligation products obtained for A7836, A11110 and A12324 were normalized to *Apc* mRNA abundance by performing SELECT with the control site A11121 (A12324: Empty, n=4; ALKBH5 OE, n=4; A11110: Empty, n=4; ALKBH5 OE, n=4; A7836: Empty, n=3; ALKBH5 OE, n=3). (D) qPCR quantification of *Apc* mRNA normalized to GAPDH obtained from Neuro2a cells transfected with shControl, sh*Ythdf1* or sh*Mettl14* (shControl, n=10; sh*Ythdf1*, n=10; sh*Mettl14*, n=9). (E) Representative images of *Apc* ViewRNA ISH (green) and YTHDF1 (purple) stainings in DIV6 neurons transfected with shControl, sh*Ythdf1* or sh*Mettl14* together with a GFP-expressing plasmid (red). Scale bar, 10µm. (F) Quantification of the number of *Apc* mRNA puncta in the cell soma (left, number of neurons : shControl, n = 24; sh*Ythdf1*, n = 21; sh*Mettl14*, n = 15) and the percentage of *Apc* mRNA in the nucleus (right, number of neurons : shControl, n = 23; sh*Ythdf1*, n = 22; sh*Mettl14*, n = 15). The graphs represent the mean ± SD (C and D) and the mean ± SEM in (F). *p<0.05, **p<0.01 or ns (not significant) comparing conditions to each other using ordinary one-way ANOVA tests (D, and F) and unpaired t-test (C).

### APC protein translation in the neuronal somata is controlled by the m^6^A machinery

Since YTHDF1 and METTL14 deregulations had no impact on *Apc* mRNA expression and export, we sought to elucidate whether m^6^A addition and reading play a role in the regulation of APC protein translation. Using a proximity ligation assay (PLA) in puromycin-treated neurons, where anti-puromycin and anti-APC antibodies were utilized, we visualized de novo protein synthesis events, as indicated by puro-PLA puncta (**Fig. 2A**) (tom Dieck et al., 2015). First, the subcellular distribution of APC translation events was assessed by imaging APC puro-PLA puncta in control cultured neurons. The puncta were detected in both soma and dendritic compartments (**Fig. 2B**). Given the crucial role of APC in the regulation of axon development, we were surprised to observe no local translation of APC in the axon/growth cone, which suggests that APC function in this region primarily relies on the control of its protein transport within the axon, rather than local translation (**Fig. 2B**). However, given the robust APC puro-PLA signal in the cytosol of the cell soma and dendrites (**Fig. 2B**), the impact of YTHDF1 and METTL14 knockdowns on APC translation in the cell soma was examined. Neurons transfected with sh*Ythdf1* and sh*Mettl14* exhibited a significant reduction in newly synthesized APC punctae, indicating the crucial role of m^6^A addition on *Apc* mRNA and its subsequent reading by YTHDF1 in proper APC protein synthesis in the neuronal cell soma and dendrites (**Fig. 2C and 2D**). To further corroborate the association between m^6^A functional loss and the APC protein translation defect, immunostaining and western blot analyses were performed on cultured neurons and Neuro2a cells, respectively. The knockdowns of YTHDF1 or METTL14 at DIV2 induce a decrease of the APC fluorescence signal in the cell soma and neurites of DIV6 neurons (**Fig. 2E, 2F and 2G**). Additionally, the knockdown of YTHDF1 in Neuro2a cells led to a reduction of APC protein level in the protein extracts as observed by western blot (**Fig. 2H and 2I**). These findings highlight the crucial role of the m^6^A machinery in facilitating proper APC protein translation and maintaining its expression in neurons.

**Figure 2.**
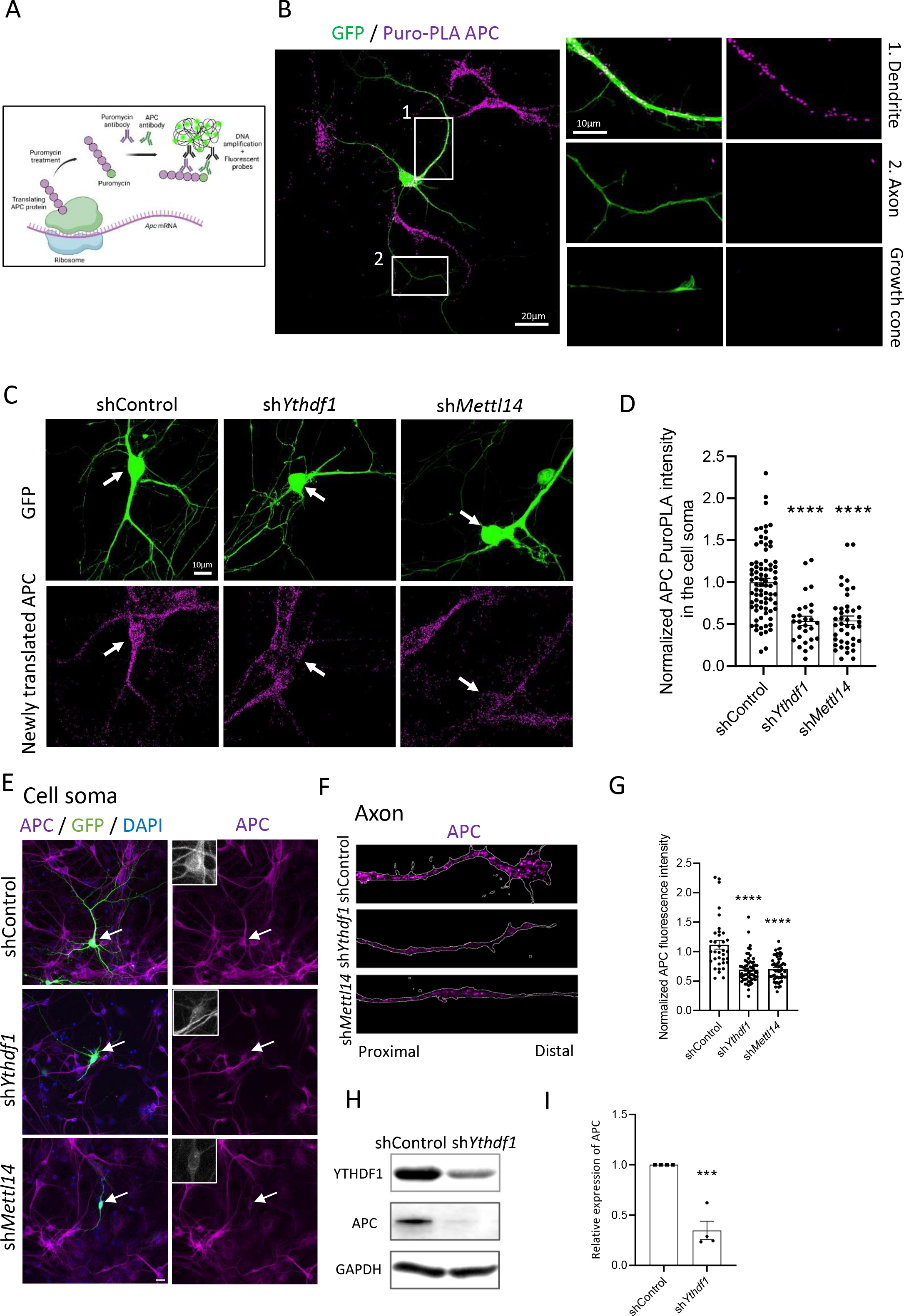
The m^6^A machinery regulates APC protein translation in the neuronal somata. A) Illustration of the puro-PLA assay used to detect the newly synthesized APC proteins. (B) Representative image of APC puro-PLA labeling (purple) in a DIV6 neuron expressing GFP (green). Scale bar, 20µm. Panels (1) and (2) provide higher magnifications of the boxed areas from the left panel located in a primary dendrite and in the axon, respectively. The right panel shows a higher magnification of the corresponding axonal growth cone. Scale bar, 10µm. (C) Representative images showing newly synthesized APC puro-PLA signal (purple) in DIV6 neurons transfected with either shControl, sh*Ythdf1* or sh*Mettl14* along with a GFP-expressing plasmid (green), treated with puromycin for 5 minutes. Transfected neurons are indicated by arrowheads. Scale bar, 10µm. (D) Quantification of APC Puro-PLA fluorescence signal intensity in the cell soma (number of cells: shControl, n=80; sh*Ythdf1*, n=28; sh*Mettl14*, n=42). (E-F) Representative images of cell soma (E) and axons (F) from neurons transfected with shControl, sh*Ythdf1* or sh*Mettl14* along with a GFP-expressing plasmid, labeled with an anti-APC antibody (magenta). (G) Quantification of APC fluorescence intensity in the cell soma and neurites of neurons transfected with shControl, sh*Ythdf1* or sh*Mettl14* (number of neurons: shControl, n=35; sh*Ythdf1*, n=62; sh*Mettl14*, n=55). Scale bar, 20µm. (H-I) Representative immunoblots (H) and quantitative analysis (I) showing the reduction of APC protein level upon YTHDF1 downregulation. GAPDH was used as a loading control for normalization (shControl, n = 4; sh*Ythdf1*, n = 4). The graphs represent the mean ± SEM in (D, G and I). ***p<0.001, ****p<0.0001 or ns (not significant) comparing conditions to each other using ordinary one-way ANOVA tests (D and G) and unpaired t-test (I).

### Downregulation of YTHDF1 disrupts microtubule and actin dynamics in the axonal growth cone

APC, known to localize in the axonal growth cones, plays a pivotal role in regulating growth cone dynamics and axon elongation (Zhou et al., 2004; Koester et al., 2007; Purro et al., 2008; Preitner et al., 2014; Efimova et al., 2020). To validate the significance of APC in axon development and growth cone dynamics, we conducted cultures of dissociated neurons and transfected them with a validated shRNA against *Apc* (**Fig. S2A**) or shControl, along with an EGFP-expressing plasmid at DIV2, and analyzed growth cone area and shape at DIV6. Downregulation of APC resulted in a decreased growth cone area, with a transition from a fan-shaped structure to a collapsed flat shape tip (**Fig. 3A and 3B**). In parallel, we performed similar analyses following knockdowns of YTHDF1 and METTL14, revealing comparable growth cone morphology defects as observed in *Apc* shRNA experiments (**Fig. 3A and 3B**). This suggests that METTL14, YTHDF1 and APC could be part of a common molecular pathway to regulate the growth cone morphology. To explore the specificity of axon defects and rule out the impact on other proteins known to be crucial for the modulation of the cytoskeleton in the growth cone, we performed immunostainings for collapsin response mediator protein 2 (CRMP2) and doublecortin (DCX). No significant changes in the immunofluorescence signal of these proteins were observed in neuronal subcompartments upon YTHDF1 downregulation (**Fig. S2B and S2C**). APC’s role in regulating the actin and microtubule cytoskeleton in the growth cone, demonstrated by its downregulation leading to decreased F-actin signal and increased microtubule growth rate in U2OS cells (Juanes et al., 2017; Efimova et al., 2020), prompted us to investigate whether similar defects would occur following YTHDF1 deregulation. Cultured neurons were labeled with markers for actin filaments (F-actin, phalloidin) and dynamic microtubules (tyrosinated tubulin). As expected, in the shControl-transfected neurons, the F-actin signal is predominantly localized to the periphery of the growth cone and organized in bundles while microtubules were primarily situated in the central domain with a few bundles invading the periphery (**Fig. 3C**). Contrastingly, YTHDF1 downregulation resulted in a significant decrease in the F-actin fluorescence signal and the size of the F-actin area in the growth cone, coupled with increased invasion of the growth cone area by dynamic microtubules, without altering fluorescence intensity (**Fig. 3C, 3D and 2E**). Further exploration of microtubule growth dynamics in the growth cone through live-imaging of EB3-NeonGreen comets, allowing tracking of microtubule growing ends, revealed a higher mean growth speed and growth length of polymerizing microtubules upon YTHDF1 downregulation compared to the control, with no impact on the duration of microtubule growth events (**Fig. 3F, 3G and 3H**). Collectively, these data demonstrate a novel role for the m^6^A machinery in governing the actin and microtubule cytoskeleton in the growth cone, which is likely to be mediated through the control of APC protein synthesis.

**Figure 3.**
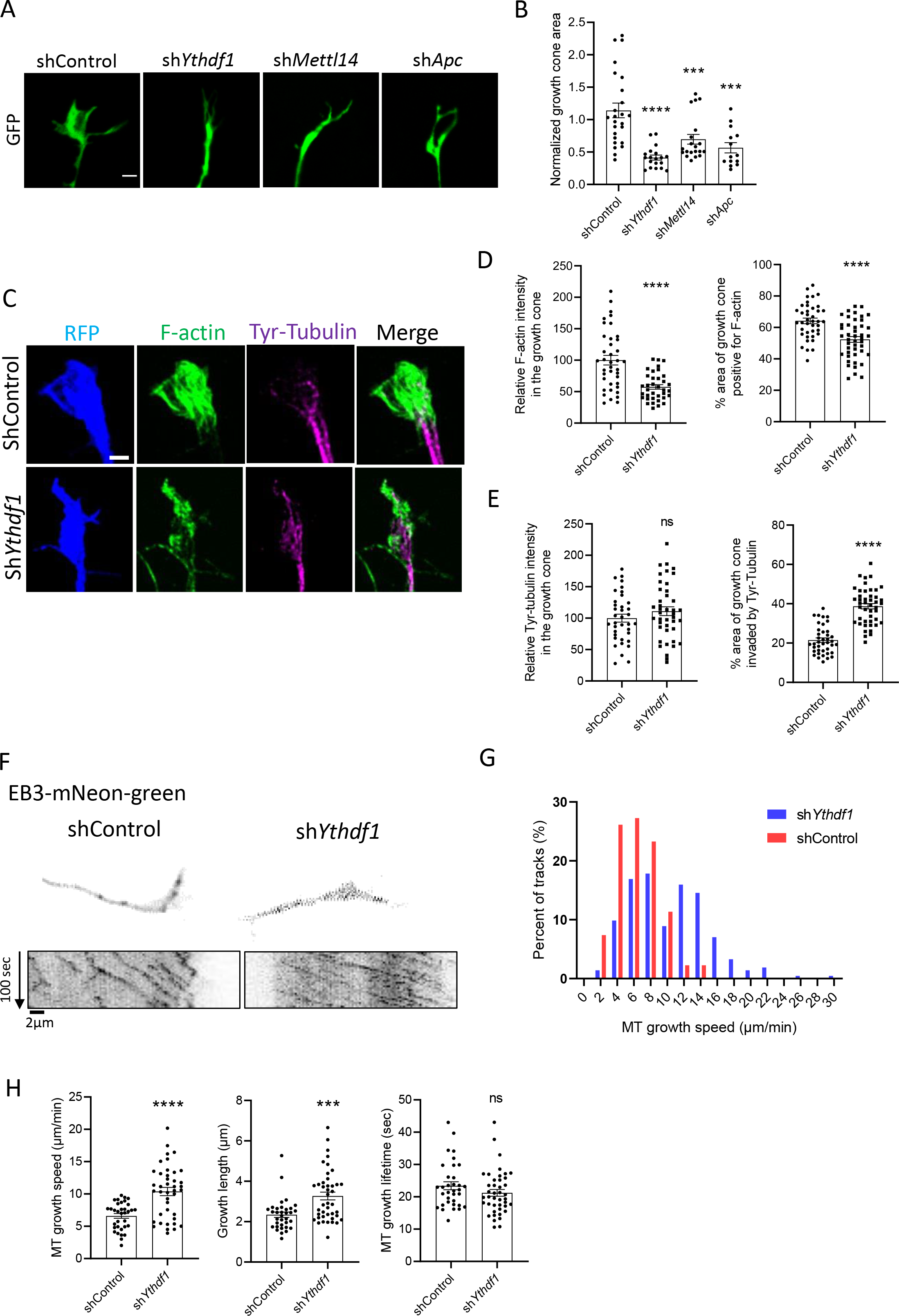
YTHDF1 downregulation induces defects in actin organization and microtubule dynamics in the axonal growth cone. (A) Representative images showing the growth cone of DIV6 neurons transfected with an EGFP-expressing plasmid and either the shControl, sh*Ythdf1,* sh*Mettl14 or* sh*Apc* at DIV2. Scale bar, 5 μm. (B) Quantification of the growth cone area (number of growth cones: shControl, n = 25; sh*Ythdf1*, n = 19; sh*Mettl14*; n=20; sh*Apc,* n=14). (C) Representative images of growth cones from DIV6 neurons expressing shControl or sh*Ythdf1* along with RFP (blue) and labeled with phalloidin (F-actin, green) and an anti-tyrosinated-tubulin antibody (Tyr-Tubulin, purple). Scale bar, 1µm. (D) Quantifications of the F-actin intensity in the growth cone (number of growth cone : shControl, n = 39; sh*Ythdf1*, n =36) and the percentage of the growth cone area positive for F-actin (number of growth cone: shControl, n = 40; sh*Ythdf1*, n = 45). (E) Quantification of the Tyr-tubulin fluorescence intensity in the growth cone (number of growth cones: shControl, n = 37; sh*Ythdf1*, n = 42) and the percentage of the growth cone area invaded by Tyr-tubulin (number of growth cones: shControl, n = 40; sh*Ythdf1*, n = 45). (F) Representative images showing the region of interest used for the live-imaging of EB3-NeonGreen comets in the growth cone of neurons transfected with shControl or sh*Ythdf1*. The kymographs generated from the region of interest are indicated below each condition. The *x*-axis corresponds to the position of the particles (µm) and the *y*-axis to the time (seconds, from top to bottom). Scale bar, 2µm. (G) Graph representing the distribution of microtubule growth speed in shControl and sh*Ythdf1* conditions. (H) Quantification of the microtubule growth speed, microtubule growth length and microtubule growth lifetime (number of comets : shControl, n=34; sh*Ythdf1*, n=42). The graphs represent the mean ± SEM in (B, D, E and H). ***p<0.001, ****p<0.0001 or ns (not significant) comparing conditions to each other using ordinary one-way ANOVA tests (B) and unpaired t-test (D, E and H).

### The m^6^A/YTHDF1/APC pathway regulates β-actin mRNA transport and local translation in the axon

To gain further insights into the molecular mechanism governing growth cone dynamics by the m^6^A machinery and APC, we focused on the regulation of β-actin mRNA transport and local translation for the following reasons: APC was shown to modulate β-actin mRNA transport by acting as a scaffold protein between molecular motors and β-actin mRNA (Baumann et al., 2020; Baumann et al., 2022); our results revealed that YTHDF1-knockdown leads to a reduction in the F-actin signal in the growth cone (**Fig. 3C, 3D**). To detect β-actin mRNA, we employed a strategy similar to the *Apc* mRNA single molecule visualization, utilizing neurons transfected with shRNA against *Ythdf1*, *Mettl14*, or the control. YTHDF1-knockdown had no effect on the total level of β-actin mRNA in the cell soma and its nuclear export, while METTL14 downregulation led to an increase in the total level of β-actin mRNA without affecting the nuclear export, which could be induced by an m^6^A-dependent destabilization mechanism of β-actin mRNA (**Fig. 4A, 4B and 4C**). Due to technical constraints in quantifying β-actin mRNA in the axon (e.g: dim GFP fluorescence in the axon after ViewRNA ISH cell assay reagent and high-temperature treatments), we performed confocal live-imaging of the fluorescently labeled β-actin mRNA through the PP7-based labeling system in the axon of DIV6 neurons (**Fig. 4D**). YTHDF1 downregulation did not alter the overall β-actin mRNA axonal transport dynamics parameters, including average speed, cumulative distance, pausing time and percentage of stationary RNA granules (**Fig. 4E and 4F**). However, we observed that YTHDF1-knockdown leads to a decrease of the β-actin mRNA granule density in the axon, indicating that less β-actin mRNA is transported in the axon in comparison to the control (**Fig. 4E and 4F**). As a decrease of β-actin mRNA in the axon could result in a defect in β-actin local translation upon YTHDF1 downregulation, we assessed the level of β-actin *de novo* protein synthesis in the different neuronal subcellular compartments using the puro-PLA method. Immunofluorescence images and quantitative analyses showed that YTHDF1-knockdown or METTL14-knockdown induces a significant decrease in the number of newly synthesized β-actin puncta in the axon shaft and in the growth cone, while β-actin protein synthesis remained unaffected in the cell soma (**Fig. 4G, 4H, S3A and S3B**). To validate that the impaired β-actin local translation in the axon is due to a m^6^A-dependent control of APC protein expression, we conducted rescue experiments in which we analyzed β-actin puro-PLA signal in the axon of primary neurons transfected with an HA-tagged APC-expressing plasmid along with either shControl, sh*Ythdf1* or sh*Mettl14.* We first validated that the expression of an HA-tagged APC protein under the control of a strong ubiquitous promoter compensates for the loss of APC protein induced by the deregulation of YTHDF1 (**Fig. S3C**). Then, we observed that the forced expression of APC in sh*Ythdf1* and sh*Mett14* transfected neurons could restore β-actin local translation in the axon and growth cone to a normal level (**Fig. 4G and 4H**). Altogether, these data suggest that the m^6^A/YTHDF1/APC pathway is required for the transport of β-actin mRNA into the axon and growth cone to ultimately regulate β-actin local protein synthesis.

**Figure 4.**
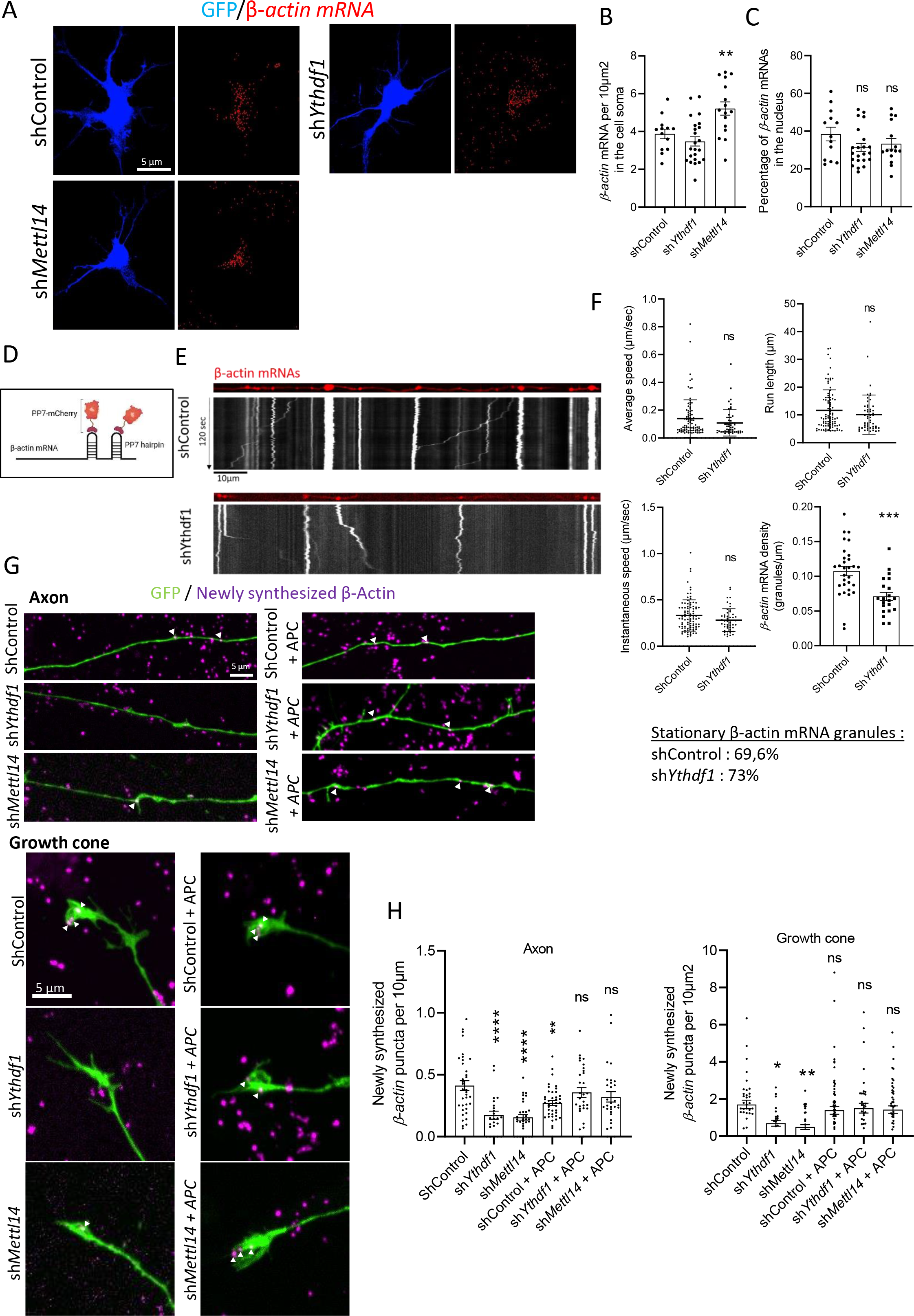
Deregulation of the m^6^A/YTHDF1/APC molecular pathway causes defects in β-actin mRNA axonal localization and local translation. (A) Representative images of *Actb* ViewRNA ISH (red) staining in DIV6 neurons transfected with shControl, sh*Ythdf1* or sh*Mettl14* along with a GFP-expressing plasmid (blue). Scale bar, 5µm. (B) Quantification of the number of β-actin mRNAs per 10µm² within the cell soma (number of cells: shControl, n=13; sh*Ythdf1*, n = 23; sh*Mettl14*; n=16). (C) Percentage of the cell soma β-actin mRNAs in the nucleus (number of cells: shControl, n=13; sh*Ythdf1*, n = 23; sh*Mettl14*; n=16). (D) Illustration of the PP7-based mRNA tagging system to track β-actin mRNA by live-imaging. (E) Representative images showing the region of interest used for the live recording of β-actin mRNA particles (red) in the axon of DIV6 neurons transfected with the β-actin mRNA reporter and either shControl or sh*Ythdf1*. The kymographs generated from the region of interest are indicated below each condition. The *x*-axis corresponds to the position of the particles (µm) and the *y*-axis to the time (seconds, from top to bottom). Scale bar, 10µm. (F) Quantification of the average speed, run length, instantaneous speed (number of granules: shControl, n=91; sh*Ythdf1*, n=52) and β-actin mRNA granule density (number of axons: shControl, n=30; sh*Ythdf1*, n=21). (G) Representative images of the Puro-PLA newly synthesized β-actin signal (purple) in the axons and growth cones of shControl, sh*Ythdf1*, sh*Mettl14*, shControl + APC-HA, sh*Ythdf1 +* APC-HA *and* sh*Mettl14 +* APC-HA DIV6 neurons. Scale bars, 5µm. (H) Quantification of the number of β-actin puro-PLA punctae in the axons and growth cones of the conditions depicted in (G) (number of growth cones: ShControl, n=37; sh*Ythdf1*, n=25; sh*Mettl14*, n=35; shControl + APC, n=65; sh*Ythdf1* + APC, n=38; sh*Mettl14* + APC, n=50; number of axons: ShControl, n=36; sh*Ythdf1*, n=23; sh*Mettl14*, n=32; shControl + APC, n=40; sh*Ythdf1* + APC, n=30; sh*Mettl14* + APC, n=30). The graphs represent the mean ± SEM in (B, C, H and F for β-actin mRNA density) and mean ± SD in (F). *p<0.05, **p<0.01, ***p<0.001, ****p<0.0001 or ns (not significant) comparing conditions to each other using ordinary one-way ANOVA tests (B, C and H) and unpaired t-test (F).

### The m^6^A/YTHDF1/APC pathway is essential for promoting axonal growth both in cultured neurons and in callosal projection neurons *in vivo*

To elucidate the necessity of the m^6^A/YTHDF1/APC pathway in axonal development, we introduced shRNAs targeting *Ythdf1*, *Mettl14*, or *Apc* mRNAs along with EGFP into DIV2 dissociated hippocampal neurons through lipofection, and subsequently fixed the cells at DIV6. In comparison to neurons transfected with shControl, those expressing shRNAs against *Ythdf1*, *Mettl14*, and *Apc* exhibited a severe impairment in axonal growth, characterized by reduced primary branch length and total axonal branches (**Fig. 5A and 5B**). As for the growth cone defects depicted in Fig. 2A-B, the analogous axonal growth defects observed upon deregulation of the m^6^A machinery and APC imply their involvement in a shared molecular pathway governing axon development. To validate that the reduction in APC protein level is accountable for the axonal defects observed upon YTHDF1 knockdown, we co-transfected neurons at DIV2 with plasmids expressing the shRNA against *Ythdf1* or the control alongside an APC-expressing plasmid and analyzed the axon length and branching at DIV6. The reexpression of APC protein in YTHDF1-knockdown neurons mitigated the axonal defects, primarily compensating for the impairment in axonal branching (**Fig. 5C and 5D**). We then sought to confirm that this function in axonal development is conserved *in vivo*. First, we assessed the expression of YTHDF1 in the mouse neonatal brain at postnatal day 1 (P1), a critical period for callosal neurons projecting axons interhemispherically through the corpus callosum. We performed immunostainings of YTHDF1, with an antibody validated on brain slices of a YTHDF1-KO mouse model (data not shown), on coronal brain slices counterstained with DAPI and we observed that YTHDF1 is widely expressed across the mouse brain, with an enrichment in the various layers of the cerebral cortex and the hippocampus (**Fig. S4A**). We then analyzed axon projections of callosal neurons on brain slices at P1 after *in utero* electroporation of tamoxifen-inducible sh*Ythdf*1 constructs in the brain lateral ventricles of embryos at embryonic day 14 (E14) (**Fig. 5E and 5F**). We performed successive injections of the pregnant mothers with tamoxifen at E17 and E18 to maintain the expression of YTHDF1 during the proliferation and migration of cortical neurons and validated that the Cre recombinase was expressed in the electroporated cortical projection neurons (**Fig. 5F and S4B**). By analyzing the length of the axons projecting to the corpus callosum, we showed that downregulation of YTHDF1 leads to a decrease in the length of the axons compared to the neurons electroporated with the shControl (**Fig. 5G and 5H**), indicating that the proper expression of YTHDF1 is required for axonal growth *in vivo*. Taken together, these data suggest that the m^6^A/YTHDF1/APC molecular pathway is indispensable for the proper development of the axon, both in cultured neurons and during cortical development in callosal projection neurons.

**Figure 5.**
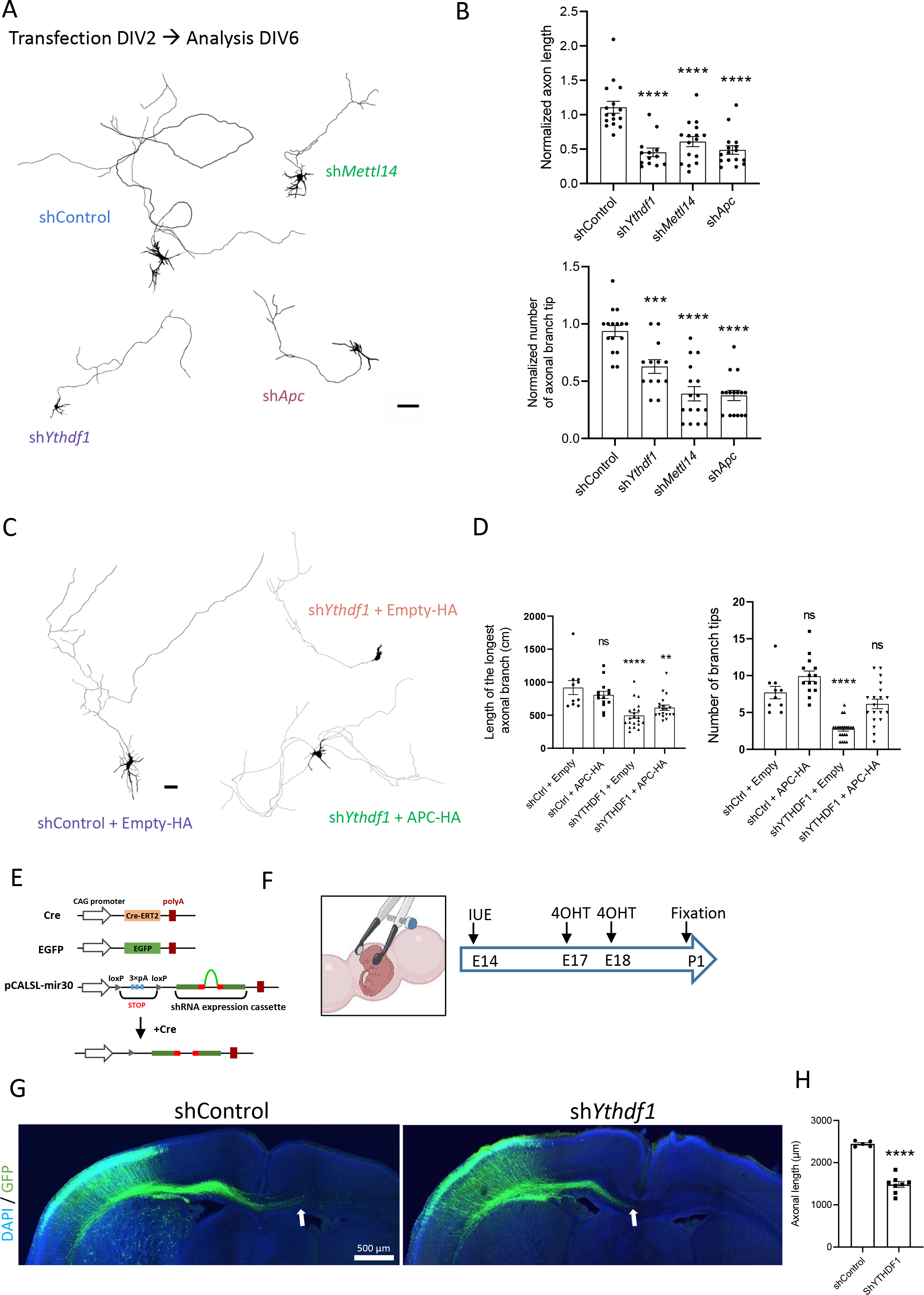
The m^6^A/YTHDF1/APC pathway is required for axonal growth in cultured neurons and callosal projection neurons *in vivo*. (A) Representative DIV6 neurons expressing either shControl, sh*Ythdf1,* sh*Mettl14 or* sh*Apc* along with EGFP. Scale bar, 50 μm. (B). Quantification of the longest axonal branch length (number of axons: shControl, n=16; sh*Ythdf1*, n=13; sh*Mettl14*; n=16; sh*Apc,* n=16) and the total number of axonal branch tips (shControl, n=16; sh*Ythdf1*, n=13; sh*Mettl14*; n=16; sh*Apc,* n=16). (C-D) Representative DIV6 neurons (C) and quantifications (D) of axon length and branching showing the partial rescue of axon growth defects by re-expressing APC in YTHDF1-knockdown neurons (number of axons: shControl + Empty, n=10; shControl + APC-HA, n=14; sh*Ythdf1 +* Empty*, n=21;* sh*Ythdf1 +* APC-HA, n=19). Scale bar, 50µm. (E) Plasmid constructs used for the Cre-inducible expression of shRNA in callosal projection neurons. (F) The experimental scheme of IUE (E14), tamoxifen (4OHT) injections (E17-E18) and brain tissue collection and fixation at P1 for imaging and analysis. (G) Representative images of P1 brain slices showing the electroporated callosal neurons expressing either shControl or sh*Ythdf1* along with EGFP and projecting contralaterally. The arrows indicate the tip of the axon bundles. Scale bar, 500µm. (H) Quantification of the maximum axonal length of electroporated callosal neurons (number of brains: shControl, n = 5; sh*Ythdf1*, n = 8). The graphs represent the mean ± SEM in (B, D and H). *p<0.05, **p<0.01, ***p<0.001, ****p<0.0001 or ns (not significant) comparing conditions to each other using ordinary one-way ANOVA tests (B and D) and unpaired t-test (H).

## Discussion

Axonal mRNA transport and local translation are crucial processes for the development of neuronal networks. One key cellular machinery regulating these processes is the interaction of RBPs with mRNAs to form RNP transport granules that can associate with molecular motors for their active transport into the axon shaft and terminals. However, our understanding of the precise mechanisms through which specific mRNAs are localized in the axonal compartment remains limited. Here, we demonstrate that the m^6^A machinery is involved in the control of the protein level of an RBP, APC, which is crucial for the regulation of the axonal cytoskeleton dynamics at the growth cone by mediating the axonal transport and local translation of β-actin mRNA (**Fig. 6**).

**Figure 6.**
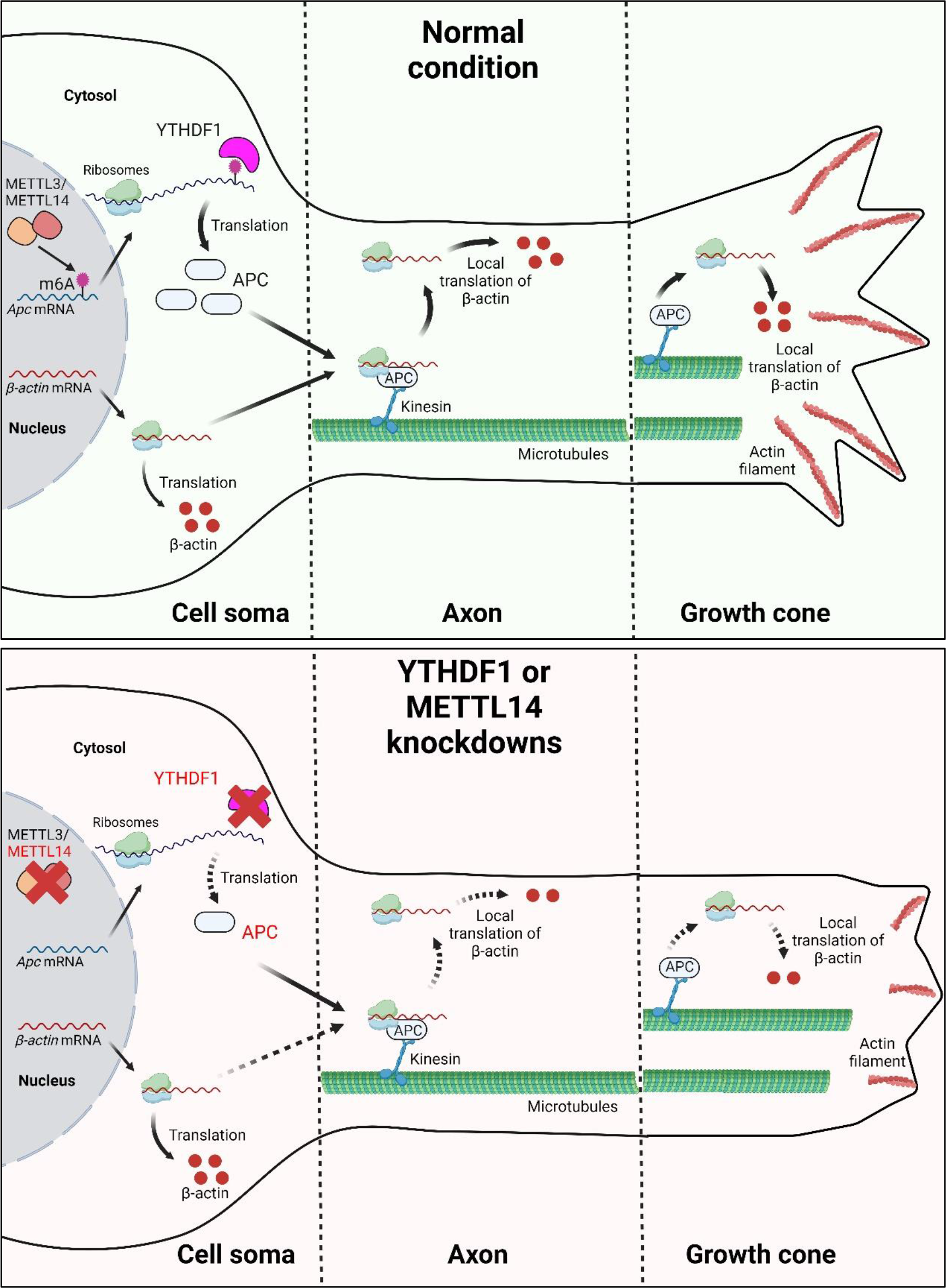
Model illustrating the regulation of axon development through the coordinated action of the m6A machinery and APC protein in controlling β-actin mRNA transport and local translation.

Two recent studies examining the neuronal subcellular localization of selected mRNAs have seemingly reached contradictory findings regarding the role of m^6^A in this process. In the first study, the authors showed that m^6^A residues in the 3′UTR promote the localization of specific neuronal mRNAs to dendrites and axons, with profiling revealing thousands of enriched m^6^A sites in neurites compared to cell bodies (Flamand and Meyer, 2022). They also show that depletion of METTL3 alters the localization of various transcripts, including *Camk2a* and *Map2*, which require m^6^A residues for neurite localization, suggesting a role for m^6^A in neuronal mRNA localization. Conversely, in the second study, the authors demonstrated that m^6^A-enriched transcripts are biased towards the cell soma versus axons, which is a consequence of lower stability (Loedige et al., 2023). Our results introduce a new model in which the m^6^A-mediated control of APC protein translation in the cell soma could regulate the transport and local translation of hundreds of mRNAs related to APC RNP granules, regardless of their m^6^A modification status. It was previously demonstrated that in the brain, APC binds to 260 mRNAs that are primarily encoding cytoskeletal proteins related to APC functions (Preitner et al., 2014). Through hypergeometric analysis, we observed a significant portion of APC-bound mRNAs (70%) intersecting with our previously published m^6^A-mRNA database (Preitner et al., 2014; Merkurjev et al., 2018), which indicates that the transport and local translation of the remaining 30% of non-m^6^A-modified APC-bound mRNAs could also be impacted by the deregulation of APC protein translation upon METTL14 or YTHDF1 deregulation. While our study focuses mainly on how the m^6^A/YTHDF1/APC pathway modulates the transport and translation of β-actin mRNA, one of APC’s target mRNAs, it would be interesting in the future to investigate whether other APC-related mRNAs are similarly deregulated and whether their m^6^A modification status is relevant. The observed disruption of microtubule dynamics in the growth cone following YTHDF1 downregulation suggests that other APC-related mRNAs such as *Tubb2b* mRNA, may also be deregulated, given APC’s role in regulating its local translation at the growth cone (Preitner et al., 2014). Furthermore, it is interesting to note that Flamand and Meyer observed that YTHDF1 depletion in neurons did not affect the transport of *CamkII* and *Map2* mRNAs (Flamand and Meyer, 2022). Given that *CamkII* and *Map2* mRNAs are not APC’s target mRNAs, these data are in accordance with our findings and suggest that YTHDF1 is involved in the specific control of the transport and translation of select mRNAs through the regulation of APC protein expression level. Is the expression of other RBPs also controlled by the m^6^A machinery through a similar mechanism? FMRP, TDP-43 and SMN1 are known to be involved in mRNA transport and local translation in neurons. Our data showed that the number of FMRP granules in axons or dendrites does not change after YTHDF1 knockdown, suggesting that m^6^A control of APC expression is not applied as a general mechanism for localized RBPs in neurons (**Fig. S5A and S5B**).

Over the past few years, there has been a growing accumulation of evidence highlighting the involvement of m^6^A signaling in regulating the development of the brain and its functions. However, despite the crucial role of the microtubule and actin cytoskeleton in maintaining neuronal functions, no connection has been established between m^6^A and the cytoskeleton to date. In our study, we revealed that YTHDF1 deregulation leads to altered microtubule and actin cytoskeleton dynamics in the growth cone and that the decrease in F-actin is likely to be due, at least partially, to a reduction in β-actin local translation. Our data indicating that APC reexpression after YTHDF1 depletion restores β-actin local translation and alleviates axon growth and branching defects further confirm the well-established role of locally synthesized β-actin in guiding axon growth and branching during neuronal development (Leung et al., 2006; Donelly et al., 2013; Yao et al., 2006; Wong et al., 2017; Bassel et al., 1998; Lepelletier et al., 2016). It is important to mention that APC was recently described as being crucial for the local assembly of branched actin filaments in the growth cone of hippocampal neurons (Efimova et al., 2020). APC targets MT tips and actin-branched regions within the growth cone to stimulate the assembly of branched-actin on MT plus-ends, and this function is likely to depend on the ability of APC to stimulate Arp2/3 signaling pathway (Efimova et al., 2020; Fang et al., 2022). It therefore remains to be delineated in future studies the specific contributions of the regulation of β-actin local translation by APC compared to its role as an MT plus-end actin branching nucleator in the maintenance of axonal growth and branching. As local translation only provides with a small amount of β-actin in comparison to the existing pool of long-lived β-actin, it is tempting to postulate that APC may additionally serve as a regulator of growth cone maintenance by coordinating local translation and the subsequent incorporation of newly synthesized β-actin at the interface between MT plus-ends and the branched-actin network.

m^6^A methylation and YTHDF readers have been implicated in the regulation of axon growth and guidance in various neuronal models and developmental stages. In particular, YTHDF1 regulates the translation of Robo3.1, a protein crucial for guiding pre-crossing axons in the embryonic spinal cord (Zhuang et al., 2019). Additionally, YTHDF1 and YTHDF2 are closely associated with axonal growth in cerebellar granule cells as the depletion of YTHDF1 and YTHDF2 enhances axonal growth in granule cells by modulating the local translation of components involved in Wnt5a signaling (Yu et al., 2021). Furthermore, in *Drosophila,* Ythdf (there is only one ortholog in *Drosophila*) plays a role in limiting axonal growth at larvae neuromuscular junctions by modulating the translation of transcripts involved in axonal growth (Worpenberg et al., 2021). Our results showing that YTHDF1 knockdown induces a reduction in axon growth and branching in primary neuron culture and in developing callosal projection neurons appear to be in contradiction with the findings in granule cells and the *Drosophila* larvae neuromuscular junction. However, our results align with a recent work demonstrating that downregulation of the long non-coding Dppa2 upstream binding RNA *(Dubr)* leads to the degradation of YTHDF1 and YTHDF3, resulting in delayed axonal growth in cortical neurons (Huang et al., 2022). Interestingly, our findings are also consistent with the previously published embryonic and postnatal axonal translatome datasets indicating that the translation of APC-related mRNAs is most pronounced in the youngest axons (E17.5) and gradually declines in postnatal stages (Shigeoka et al., 2016). This implies that YTHDF1 likely plays a critical role in regulating axonal growth during axonal development through the modulation of APC translation but that this function may vary between young and adult axons, neuronal populations and species.

Finally, given that *APC* has been identified as a risk gene for autism spectrum disorder and intellectual disabilities (Barber et al., 1994; Raedle et al., 2001; Cui et al., 2005; Finch et al., 2005; Heald et al., 2007; Zhou et al., 2007; Mohn et al., 2014), our findings suggest that deregulation of the YTHDF1/m^6^A/APC axis pathway could contribute to the physiopathology of these neurological disorders. The implication of m^6^A in neurological disorders provides novel avenues for potential therapeutic targets in treatment. Notably, m^6^A editing toolkits using the catalytically inactive CRISPR–Cas13 system for installing, removing or directing the readers to specific mRNAs have recently emerged as a promising therapeutic strategy (Tang et al., 2021; Sokpor et al., 2021). Furthermore, our findings give new insights into the regulation of local translation in neurons, which is particularly crucial for a better understanding of the pathological mechanisms underlying neurological diseases associated with defects in local translation such as neurodevelopmental disorders (Fragile X syndrome) and neurodegenerative diseases (e.g. amyotrophic lateral sclerosis, frontotemporal dementia, spinal muscular atrophy).

## Author contributions

L.B, R.R, and D.O.W conceived and designed the project. L.B designed, performed, and analyzed most experiments and wrote the manuscript. R.R performed part of the experiments and contributed to manuscript writing. I. O and H. U performed part of the experiments. D.O.W edited the manuscript.

## Acknowledgments

L.B postdoctoral fellowship was supported by the Japan Society for the Promotion of Science (JSPS). R.R was supported by the Sasakawa research grant, Mitsubishi scholarship and JASSO Honor Fellowship. This work is supported by KAKENHI (22KF0399, 21H02580) to D.O.W. Figure 6 was created with BioRender.com.

## Materials and methods

### Animal

ICR mice were purchased (SLC or Shimizu, Japan) and housed at the Institute for integrated cell-material Sciences (iCeMS) or RIKEN Center for Biosystems Dynamics Research (BDR) animal facilities. This study was carried out in accordance with the Guide for the Care and Use of Laboratory Animals from the Society for Neuroscience and was authorized by the Animal Care and Use Committee of Kyoto University and the RIKEN BDR.

### Plasmids and Oligos

For the construction of knockdown vectors, synthetic oligos were designed and cloned into BglII/HindIII sites or BglII/XhoI sites on pSUPER vector protocol (Oligoengine), and into XhoI/EcoRI sites of pCALSL-mir30 (Addgene plasmid #13786) (Matsuda & Cepko, 2007). The oligos for the knockdown experiments mentioned below are designed against mouse genome: shControl: 5’-GCTACCTCCATTTAGTGT-3’; sh*Ythdf1*: 5’-CTAATACATCCCAAAGATA-3’; sh*Mettl14*: 5’-CTTACACTTGCATCTTTAT-3’; sh*Apc*: 5’-GTACTAACTAGGTAAATA-3’. pCAG-ERT2CreERT2 plasmid was purchased from Addgene (plasmid #13777). For SELECT experiments, mouse full-length cDNA of ALKBH5 was subcloned into a pCAGGS-HA vector. For β-actin mRNA transport analysis, pHR-tdPP7-3xmCherry and pcDNA4TO-24xGCN4_v4-kif18b-24xPP7 were purchased from Addgene (plasmids #74926 and #74928, respectively). Briefly, the CMV promoter was replaced by a CAG promoter using HindIII/MluI and the mus musculus β-actin mRNA CDS (NM_007393.5) was inserted using NheI/AgeI. The 5’UTR was inserted using HindIII/HindIII and finally, the 3’UTR was inserted using BamHI/NheI to obtain the reporter plasmid 5’UTR-SunTag24x-Actin-3’UTR-PP724x. pCAG-EGFP plasmid was a kind gift from Kengaku Mineko’s lab (iCeMS, Kyoto University, Japan). Mus musculus *Apc* cDNA (NM_001402731.1) was subcloned in a pCAGGS-HA vector using BamHI/EcoRI.

### Neuronal cell culture and transfection

Hippocampi were dissected from postnatal day 0 (P0) mice. Hippocampal neurons were dissociated using neuron dissociation solutions (Wako) and plated on 12 mm coverslips or 35 mm glass-bottom dishes (MatTek) coated with poly-L-lysine (Sigma) in MEM (Gibco) supplemented with 10% horse serum (Gibco), 0.6% D-glucose (Sigma), 1 mM sodium pyruvate (Nacalai Tesque) and 1% penicillin-streptomycin (Gibco). Three hours after plating, the media was replaced by growth media consisting of Neurobasal-A medium (Gibco) supplemented with B-27 supplement, GlutaMAX and penicillin-streptomycin. Cells were maintained at 37°C, 5% CO_2_ until fixation or live-imaging.

### Immunostaining in primary neuronal culture

Dissociated hippocampal cultures (DIV6) were fixed in 4% PFA/PBS and permeabilized in 0.1% tritonX-100/PBS for 10 min at RT. Cells were blocked in 10% donkey serum/PBS for at least 1 hour before primary antibody incubation (in 10% donkey serum/PBS, overnight, 4°C). Next day, after 3 times washing with 1X PBS, followed by secondary antibody incubation in the dark for 1.5 hours, samples were counterstained with DAPI (1 µg/ml in PBS for 10 min at RT) and mounted onto glass slides with ProLong™ Diamond Antifade Mountant (ThermoFisher). To label F-actin in axonal growth cones, dissociated hippocampal cultures (DIV6) were fixed in 4% PFA (20 mins at 37°C), 3 times washed with 1X PBS and permeabilized in 0.1% triton X-100/PBS for 10 min at RT. For single staining, Phalloidin conjugated with fluorophore (488 or 594) solution was diluted at 1:1000 to 1:500 using the dilution buffer composed of 1X PBS and 1% BSA. The neurons with the Phalloidin solution were incubated for 90 minutes. After 3 times washing with 1X PBS, followed by MilliQ wash, samples were mounted onto glass slides with ProLong™ Diamond Antifade Mountant (ThermoFisher). Images of tyrosinated-tubulin and F-actin stainings in the growth cone were acquired using a confocal microscope system Dragonfly 500 (Oxford Instruments).

### Puromycin-Proximity Ligation Assay (Puro-PLA)

Puro-PLA experiments were performed as previously described (Dieck et al., 2015) to label newly synthesized β-actin and APC proteins. Briefly, neurons cultured 6 days *in vitro* were incubated with 10µg/mL puromycin for 5min, washed with warm PBS and fixed for 10 min in 4% PFA-sucrose. Following fixation and permeabilization, the Duolink assay (DUO92013) was performed as recommended by the manufacturer using the following primary antibodies: Puromycin (Merck Millipore, MABE343, 1:400), APC (Santa Cruz, sc-896, 1:200), β-actin (Cayman Chemicals, 32127, 1:200). Cells were mounted in ProLong™ Diamond Antifade Mountant (ThermoFisher) and images were acquired with a Dragonfly confocal microscope (Oxford Instruments).

### RNA fluorescence in situ hybridization (FISH) on cultured mouse neurons

DIV6 cultured neurons were fixed and processed using the ViewRNA Cell Plus Assay kit (ThermoFisher) based on the manufacturer’s instructions. Custom-made ViewRNA Cell Plus Probe Sets (ThermoFisher) against mouse *Actb* mRNA (VB6-12823-VC) and mouse *Apc* mRNA (VB6-19724-VC) were used. YTHDF1 staining was performed during the ViewRNA Cell Plus Assay experiment according to the manufacturer’s protocol. Neurons were then counterstained with DAPI and observed under a Dragonfly confocal microscope (Oxford Instruments).

### EB3 comets analysis *in vitro*

DIV2 neurons cultured in 35 mm glass-bottom dishes (MatTek) were transfected with EB3-GFP plasmid and either shControl and sh*Ythdf1* plasmids and imaging was performed at DIV6. During imaging, the dishes were maintained in the incubation chamber of the LSM800 laser scanning confocal microscope at 37°C with a continuous supply of 5% CO_2_. Images were acquired with 63X Plan Apochromat objective using ZEN software (Zeiss). Axonal growth cone images were captured every 3 seconds during 100 seconds. Kymographs were generated using the ImageJ plugin KymoToolBox.

### Live-imaging of β-actin mRNA

DIV2 neurons cultured in 35 mm glass-bottom dishes (MatTek) were transfected with the reporter construct (5’UTR-SunTag24x-Actin-3’UTR-PP724x), PP7-mCherry plasmids and either shControl and sh*Ythdf1* plasmids and imaging was performed at DIV6. During imaging, the dishes were maintained in the incubation chamber of the LSM800 laser scanning confocal microscope at 37°C with a continuous supply of 5% CO_2_. Axon images were captured at 500 ms intervals for 120 sec under a Dragonfly confocal microscope (100X magnification, Oxford Instruments). Kymographs were generated using the ImageJ plugin KymoToolBox. β-actin mRNA granules were considered as stationary if the directional transport observed was less than 4μm within 120 sec.

### Neuro2A cell culture and transfection

Mouse Neuro-2a neuroblastoma cells (ATCC® CCL-131™) were cultured in Minimum Essential Media (MEM, 1X, Gibco) supplemented with 10% (v/v) fetal bovine serum (FBS, heat-inactivated, Biosera) without any antibiotic supplement. Neuro-2a cells were maintained at 37°C in 5% CO_2_ until ready for downstream experiments. Neuro-2a cells were placed in MEM with 10% FBS without Pen-Strep 24h before transfection using Lipofectamine 2000 (Invitrogen). Media was changed 24 hours after transfection with 10% Fetal Bovine Serum containing MEM media with Pen-Strep. Neuro-2a cells were subjected to differentiation 48h after transfection using MEM media containing low serum (0.5% FBS) with ATRA (final concentration 10 µM). Cells were maintained for one more day before harvesting for western blotting or qPCR analysis.

### Western blotting

Neuro2a cell lysates were prepared using RIPA buffer after washing with ice-cold PBS. RIPA buffer composition was as follows: 50 Mm Tris-Cl pH 7.6, 150 mM NaCl, 1 mM EDTA (pH 8.0), 1% TritonX-100, 4% SDS and Ultrapure water. For 10 ml of RIPA buffer, 1 tablet of the protease inhibitor cocktail from Roche was added. The cell suspension was sonicated using TOMY Ultrasonic Disruptor UD-201. The suspension was centrifuged at 15000 RPM for 5 mins at 4℃ using a HIMAC CT15RE benchtop centrifuge. The supernatant was collected in a low-binding Eppendorf tube and further subjected to BCA assay using Pierce™ BCA Protein Assay Kit (ThermoFisher Scientific) according to the manufacturer’s protocol. Subsequently, lysate samples were boiled with 5X SDS at 95°C and 20 µg of lysates were loaded on 10% SDS– polyacrylamide gel to be finally transferred to a PVDF membrane for immunoblotting. To develop HRP chemiluminescence signal Amersham™ ECL Select™ Western Blotting Detection Reagent (Cytiva) was used in 1:1 dilution. Membranes were imaged using BioRAD GelDoc™ XR Molecular Imager. Band densitometry was performed using ImageJ.

### qPCR analysis

Transfected Neuro-2a cells were processed for RNA extraction using RNeasy Mini Kit (Qiagen) following the manufacturer’s instructions. 1µg of RNA was reverse-transcribed with iScript RT Supermix (Bio-Rad). Following this, appropriate dilution of the cDNA was then used in qPCR reactions containing PowerUp SYBR Green Master Mix (Applied Biosystems) and primer pairs. qPCR reactions were performed using the StepOnePlus Real-Time PCR System (Applied Biosystems). Analysis was done using the 2-ΔΔCT method after defining primer efficiencies, *Gapdh* served for normalization. The following primers were used: *Ythdf1*, GCATCAGAAGGATGCAGTTCATG (forward) and GATGGTGGATAGTAACTGGACAG (reverse); *Mettl14,* AGAGTGCGGATAGCATTGGTGC (forward) and CTCCTTCATCCAGACACTTCCG (reverse); *Apc,* CCCCAGTGACCTTCCAGATA (forward) and GCACTGTCTGTGGAGGAGGT (reverse).

### SELECT assay for single-base m^6^A detection

m^6^A coordinates on *Apc* mRNA were selected from various single nucleotide m^6^A sequencing studies like DART-seq (Meyer, 2019), m^6^A CLIP-seq using mouse brain samples (Ke et al., 2015; Shi et al., 2018), YTHDF1 binding site analysis (Shi et al, 2018) and our own m^6^A iCLIP dataset from mouse forebrain. All of these sites were also corroborated in m^6^A Atlas (Tang *et al*, 2021). To assess m^6^A modification levels at the selected individual sites, we designed four probe pairs specifically targeting the A7836, A11110, A11121 and A12324 sites within the *Apc* transcript (NM_007462.3) for SELECT assays. The SELECT method was performed as described previously (Xiao et al., 2018). In brief, 2.2µg of total RNA from Neuro2a cells was combined with 40 nM Up Primer, 40 nM Down Primer, and 5 μM dNTP in a 17μl solution of 1×CutSmart buffer (containing 50 mM KAc, 20 mM Tris-HAc, 10 mM MgAc2, 100 μg/ml BSA, pH 7.9 at 25°C). The RNA and primers were then annealed by subjecting the mixture to a temperature gradient: 90°C for 1min, 80°C for 1min, 70°C for 1min, 60°C for 1min, 50°C for 1min, and finally 40°C for 6min. Following annealing, a 3 μl mixture comprising 0.01 U Bst 2.0 DNA polymerase, 0.5 U SplintR ligase, and 10 nmol ATP was added to the previous mixture, bringing the final volume to 20 μl. The reaction mixture was then incubated at 40°C for 20 min, denatured at 80°C for 20 min, and finally kept at 4°C. Following this, quantitative qPCR reactions were conducted using the StepOnePlus Real-Time PCR System (Applied Biosystems). The 20μl qPCR reaction mixture comprised 2x PowerUp™ SYBR™ Green Master Mix (Applied Biosystems), 200 nM of qPCR-F primer, 200 nM of qPCR-R primer, 2μl of the final reaction mixture, and ddH2O. The qPCR protocol was performed using the following steps: 95°C for 5 minutes; (95°C for 10 seconds, 60°C for 35 seconds) repeated for 40 cycles; 95°C for 15 seconds, 60°C for 1 minute; 95°C for 15 seconds; and finally, holding at 4°C. Data analysis was performed using StepOne Real-Time PCR Software v2.3. List of SELECT oligos are mentioned in Table 1.

### *In utero* electroporation

Timed-pregnant (E14) mice were anesthetized with isoflurane before the injection of painkillers. Endotoxin-free plasmids mixed with 0.1% fast green (Sigma) were injected into lateral ventricles of the embryo forebrains, followed by electroporation (5 pulses of 40 mV at 50 ms intervals for 950 ms) using 3mm diameter platinum electrodes (Sonidel) connected to an electroporator (ECM 830, BTX). The electroporation was performed with a combination of three plasmids consisting of a tamoxifen-inducible CreERT2 expressing plasmid and a pCAGGS-EGFP plasmid along with either shControl or sh*Ythdf1*. The embryos were placed back in the abdominal cavity. For injection, 4OHT and progesterone were dissolved in 100% ethanol at concentrations of 20 mg/ml and 10 mg/ml, respectively, and then further diluted with nine volumes of corn oil (Sigma-Aldrich). The diluted tamoxifen solution (2 mg/ml, 100 µl per mouse) was administered intraperitoneally two times at E17 and E18 and the brains of the pups were dissected at P1 and fixed in 4% PFA overnight. Fixed brains were embedded in agarose 4% low melting agarose (Bio-Rad) and cut into coronal sections (100 µm) using a vibratome (Leica VT1000S, Leica Microsystems).

For immunostainings, fixed P1 brain coronal sections were permeabilized in 0.3% Triton X-100/PBS, and blocked in 10% donkey serum in 1X PBS for 1h. Primary antibodies were incubated overnight at 4°C with 0.3% Triton X-100/PBS and 1% donkey serum. The following antibodies were used: anti-Cre Recombinase (MAB3120, 1:5000, Millipore); anti-YTHDF1 (17479-1-AP, 1:200, Proteintech); anti-GFP (ab13970, 1:2000, Abcam). After washing in 0.3% Triton X-100/PBS, the sections were incubated with corresponding Alexa Fluor conjugated secondary antibodies (Invitrogen) in a 1:1000 dilution in 1X PBS for 1h30 at room temperature. After washing, sections were counterstained with DAPI and mounted on glass slides with ProLong™ Gold Antifade Mountant (ThermoFisher).

### Analysis of neuronal morphology

Fluorescence images of DIV6 neurons to study axonal growth were acquired on a BZ-X700 microscope (Nikon). Multichannel images were acquired to cover the entire territory of single neurons and stitched in the BZ-X viewer to capture the entire axon length. The Segmented Line selection tool in ImageJ software was used for tracing axons and quantifying the axon length. Two parameters were measured to assess axonal growth: 1. length of the longest axon. 2. The number of parallel branches (by counting the number of the endpoints of the axonal branches, excluding thin and short protrusions). Representative neurons were traced using the Simple Neurite Tracer (SNT) plug-in (ImageJ), and processed in Paint3D and vectorized for visualization.

### Statistical Analysis

To perform all the statistical analysis GraphPad Prism version 8 was used. The n number of each experiment and a detailed description of the method of statistical analysis and the results are mentioned in the legend of each figure. Comparison between two groups with normally distributed samples was performed with a two-sided unpaired t-test, while comparisons among more than two groups with one-way analysis of variance (ANOVA) followed by the appropriate post hoc test. The statistical notations in the graphs are as follows: ns, not significant; *p ≤ 0.05; **p ≤ 0.01; ***p ≤ 0.001; ****p ≤ 0.0001. p-value > 0.05 is considered non-significant.

