## Supplementary figures for "m^6^A RNA methylation-dependent APC translation is required for local translation of beta-actin and axon development"

**Supplementary figure S1. Validation of the efficiency of *Ythdf1* and *Mettl14* shRNAs.** (A) Quantification of the YTHDF1 fluorescence intensity in DIV6 neurons transfected at DIV2 with shControl or sh*Ythdf1* (number of cells: shControl, n=17; sh*Ythdf1*, n=15). (B) qPCR quantification of *Ythdf1* mRNA normalized to GAPDH obtained from Neuro2a cells 48h after transfection with shControl or sh*Ythdf1* (shControl, n=8; sh*Ythdf1*, n=8). (C) Quantification of the METTL14 fluorescence intensity in DIV6 neurons transfected at DIV2 with shControl or sh*Mettl14* (number of cells: shControl, n=81; sh*Mettl14*, n=35). (D) qPCR quantification of *Mettl14* mRNA normalized to GAPDH obtained from Neuro2a cells 48h after transfection with shControl or sh*Mettl14* (shControl, n=6; sh*Mettl14*, n=6). The graph represents the mean ± SEM in (A and C) and the mean ± SD in (B and D). ***p<0.001, ****p<0.0001 or ns (not significant) comparing conditions to each other using unpaired t-test.

**Supplementary figure S2. YTHDF1 knockdown does not affect DCX and CRMP2 protein levels.** (A) Validation of the sh*Apc* efficiency by quantification of APC fluorescence intensity in DIV6 neurons transfected at DIV2 with the shControl or sh*Apc* (number of cells: shControl, n=21; sh*Apc*, n=17). (B) Representative images of DIV6 neurons transfected with GFP (green) and the shControl or sh*Ythdf1* and labeled with a DCX antibody (magenta). The left part shows representative neuronal somata and the right part shows representative growth cones for both conditions. The graphs show the quantification of DCX fluorescence intensities in the cell soma (number of cells: shControl, n=15; sh*Ythdf1*, n=12) and in the growth cone (number of growth cones: shControl, n=9; sh*Ythdf1*, n=8) after expression of the shControl or sh*Ythdf1.* Scale bar, 10µm (cell soma) and 1µm (growth cone). (C) Representative images of DIV6 neurons transfected with GFP (green) and the shControl or sh*Ythdf1* and labeled with a CRMP2 antibody (magenta). The left part shows representative neuronal somata and the right part shows representative growth cones for both conditions. The graphs show the quantification of CRMP2 fluorescence intensities in the cell soma (number of cells: shControl, n=17; sh*Ythdf1*, n=16) and the growth cone (number of growth cones: shControl, n=7; sh*Ythdf1*, n=7) after expression of the shControl or sh*Ythdf1.* Scale bar, 10µm (cell soma) and 1µm (growth cone). The graphs represent the mean ± SEM in (A, B and C). ns (not significant) or ****p<0.0001 comparing conditions to each other using unpaired t-tests in (A, B and C).

**Supplementary figure S3. Quantification of Puro-PLA β-actin in the cell soma and validation of the expression of HA-tagged APC protein upon YTHDF1 knockdown.** (A) Representative images of the Puro-PLA newly synthesized β-actin signal (purple) in the cell soma of shControl, sh*Ythdf1*, sh*Mettl14* DIV6 neurons. Scale bars, 10µm. (B) Quantification of the number of β-actin puro-PLA punctae in the cell soma of the conditions depicted in (A) (number of cells: shControl, n=19; sh*Ythdf1*, n = 22; sh*Mettl14*; n=14). The graph represents the mean ± SEM. ns (not significant) comparing conditions to each other using ordinary oneway ANOVA tests. (C) Immunofluorescence stainings of APC (magenta) and HA (green) of DIV6 neurons transfected at DIV2 with GFP (blue) and either a combination of sh*Ythdf1* and an HA-tagged APC expressing vector or sh*Ythdf1* and the control empty vector. Scale bar, 10µm.

**Supplementary figure S4. YTHDF1 is expressed in the mouse neonatal brain and the Cre recombinase is expressed in electroporated callosal projection neurons.** (A) Immunostainings on P1 coronal mouse brain slices using a YTHDF1 antibody (red) and counterstained with DAPI (blue). The lower panels show higher magnifications in the somatosensory cortex (SSC) and the hippocampus (Hipp). Scale bar, upper panels 500µm; lower panels 200µm. (B) Validation of Cre recombinase conditional expression in electroporated callosal projection neurons. *In utero* electroporation was performed at E14 and the pregnant mother was injected with tamoxifen at E17 and E18. The brains were harvested at P1 and brain sections were labeled with a Cre antibody (red) and counterstained with DAPI (blue). Scale bar, 100µm.

**Supplementary figure S5. YTHDF1 knockdown does not impact the number of FMRP granules in neurites.** (A) Representative images of DIV6 axons of neurons transfected with GFP (green) and the shControl or sh*Ythdf1* and labeled with an FMRP antibody (magenta) and quantification of the number of FMRP granules per 10µm in the axon after expression of the shControl or sh*Ythdf1* (number of axons: shControl, n=11; shYthdf1, n=12). Scale bar, 5µm. (B) Representative images of DIV6 dendrites of neurons transfected with GFP (green) and the shControl or sh*Ythdf1* and immunostained with an FMRP antibody (magenta) and the bar graph showing the quantification of the number of FMRP granules per 10µm in the dendrites of shControl or sh*Ythdf1* transfected neurons (number of cells: shControl, n=11; shYthdf1, n=16). Scale bar, 5µm. The graphs represent the mean ± SEM in (A and B). ns (not significant) comparing conditions to each other using unpaired t-tests (A and B).

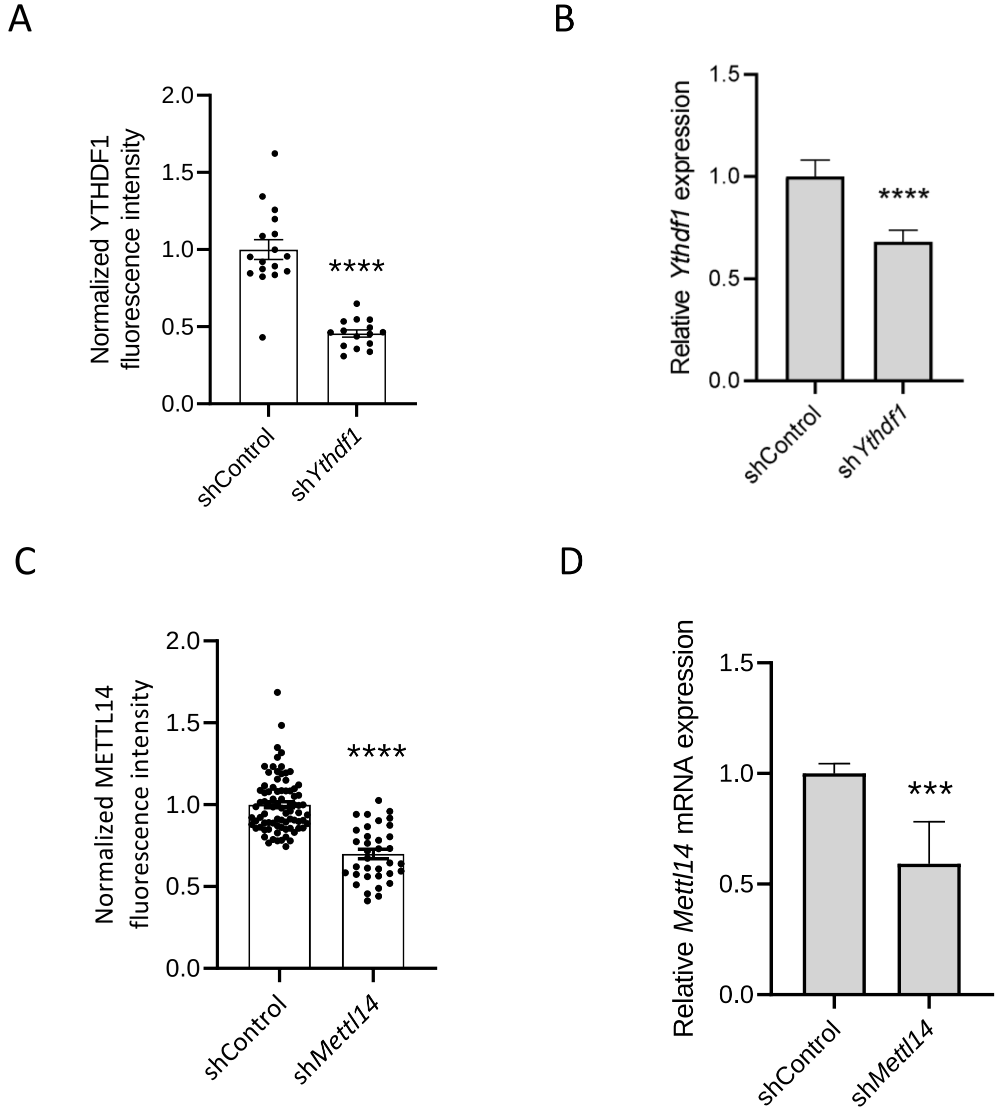
A
Normalized YTHDF1 fluorescence intensity
2.0
1.5
1.0
0.5
0.0
****
shControl
shYthdf1
B
Relative Ythdf1 expression
1.5
1.0
0.5
0.0
****
shControl
shYthdf1
C
Normalized METTL14 fluorescence intensity
2.0
1.5
1.0
0.5
0.0
****
shControl
shMettl14
D
Relative Mettl14 mRNA expression
1.5
1.0
0.5
0.0
***
shControl
shMettl14

Fig. S1

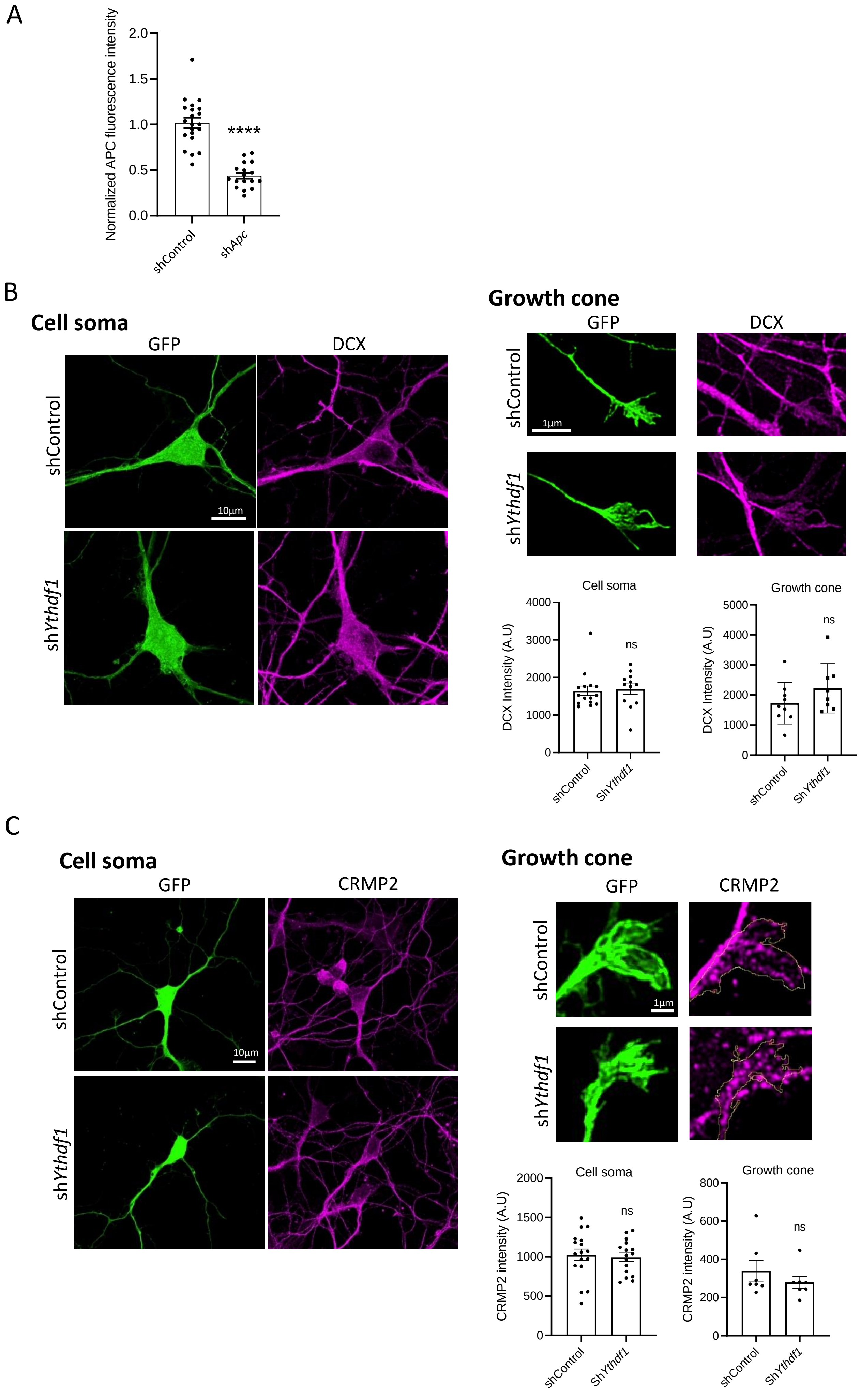
A
Normalized APC fluorescence intensity
2.0
1.5
1.0
0.5
0.0
****
shControl
shApc
B
Cell soma
GFP
DCX
shControl
10μm
shYthdf1
Growth cone
GFP
DCX
shControl
1μm
shYthdf1
Cell soma
DCX Intensity (A.U)
4000
3000
2000
1000
0
ns
shControl
ShYthdf1
Growth cone
DCX Intensity (A.U)
5000
4000
3000
2000
1000
0
ns
shControl
ShYthdf1
C
Cell soma
GFP
CRMP2
shControl
10μm
shYthdf1
Growth cone
GFP
CRMP2
shControl
1μm
shYthdf1
Cell soma
CRMP2 intensity (A.U)
2000
1500
1000
500
0
ns
shControl
ShYthdf1
Growth cone
CRMP2 intensity (A.U)
800
600
400
200
0
ns
shControl
ShYthdf1

Fig. S2

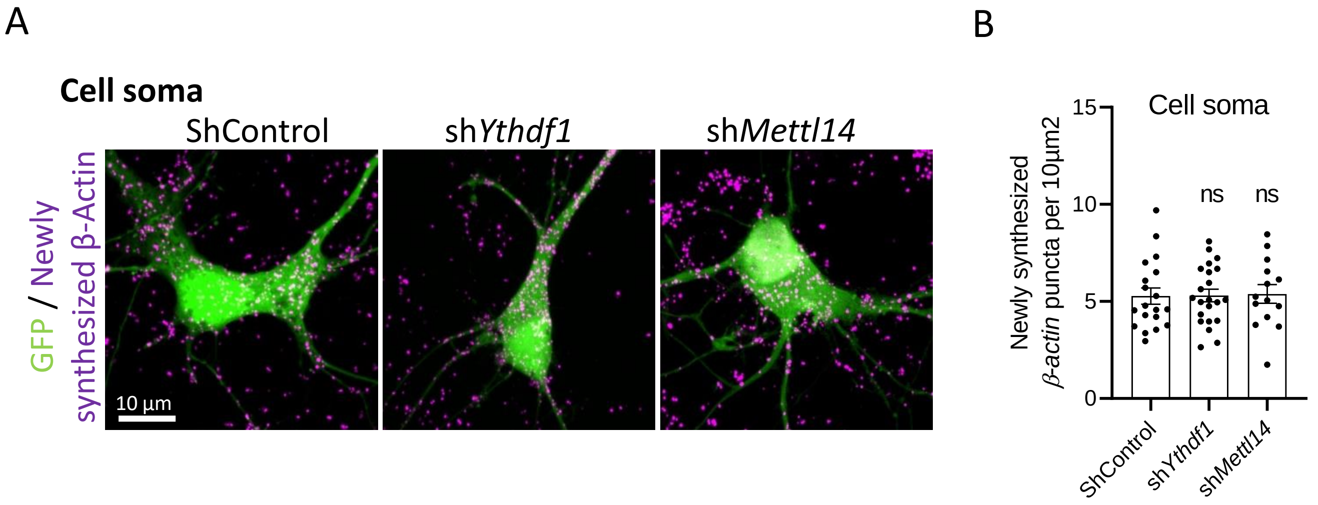
A
Cell soma
ShControl
shYthdf1
shMettl14
GFP / Newly synthesized β-Actin
10 µm
B
Cell soma
Newly synthesized β-actin puncta per 10µm2
15
10
5
0
ns
ns
ShControl
shYthdf1
shMettl14

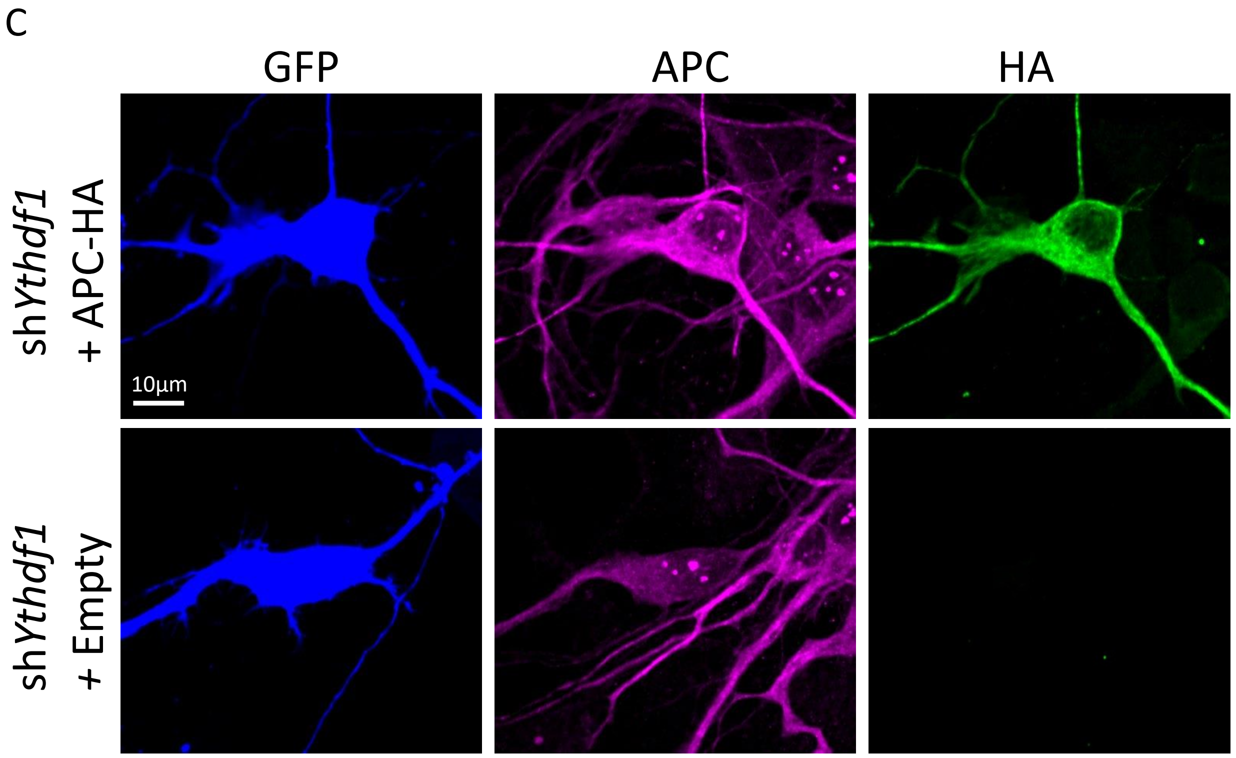
C
GFP
APC
HA
shYthdf1 + APC-HA
shYthdf1 + Empty
10µm

Fig. S3

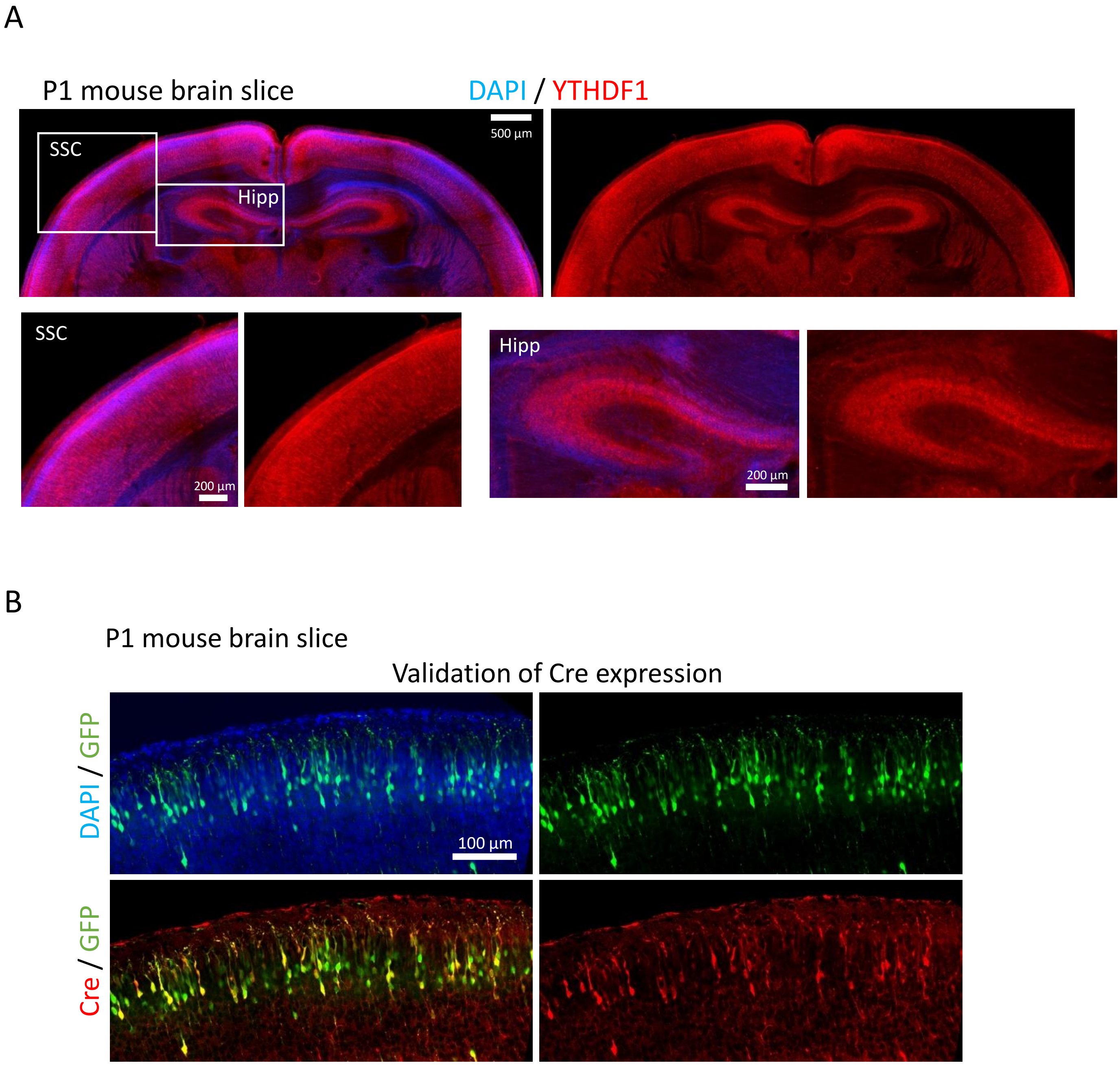
A
P1 mouse brain slice
DAPI / YTHDF1
SSC
Hipp
500 µm
SSC
200 µm
Hipp
200 µm
B
P1 mouse brain slice
Validation of Cre expression
DAPI / GFP
100 µm
Cre / GFP

Fig. S4

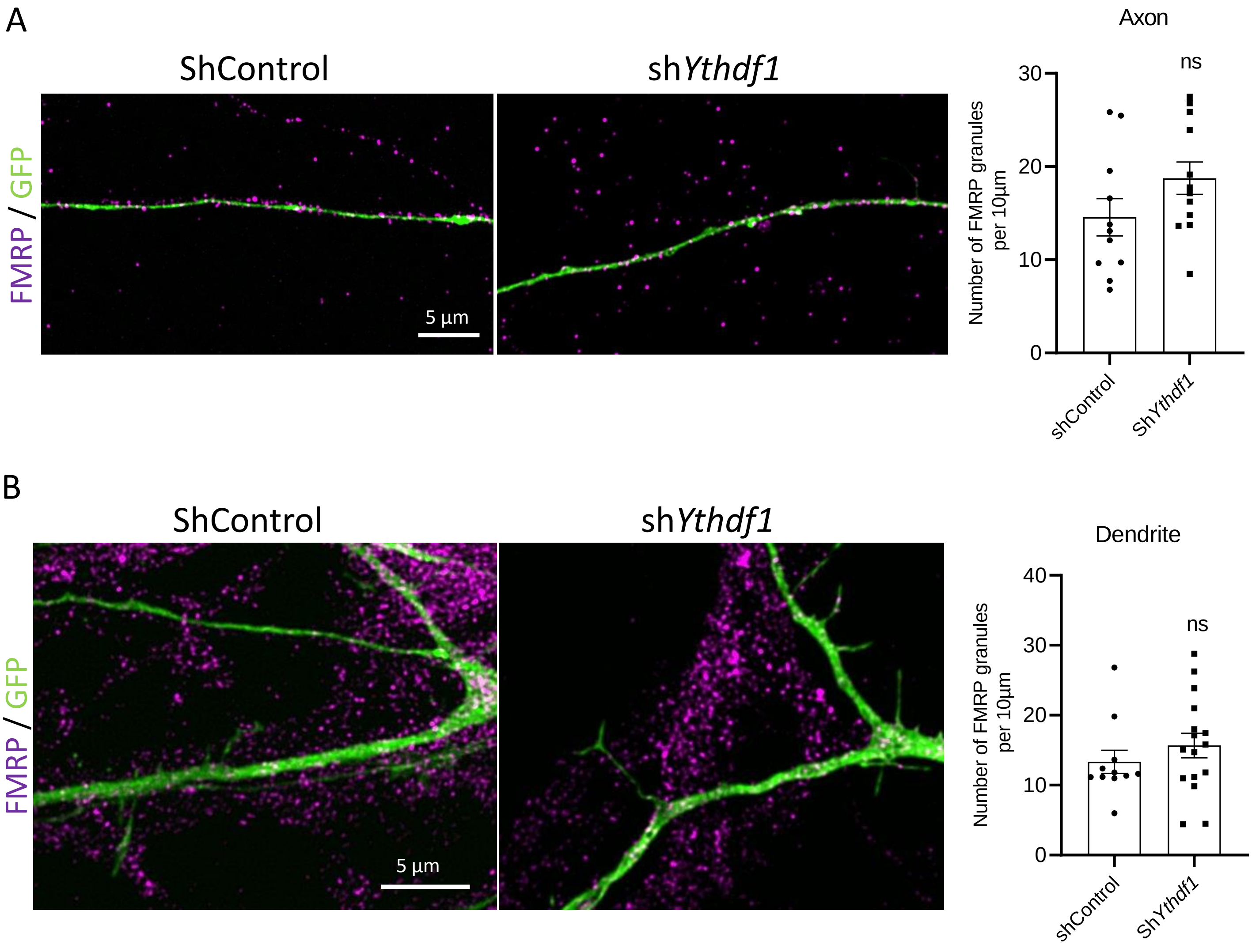
A
ShControl
shYthdf1
FMRP / GFP
5 µm
Axon
ns
30
20
10
0
Number of FMRP granules per 10µm
shControl
ShYthdf1
B
ShControl
shYthdf1
FMRP / GFP
5 µm
Dendrite
ns
40
30
20
10
0
Number of FMRP granules per 10µm
shControl
ShYthdf1

Fig. S5
